# Tyrosinase-induced neuromelanin accumulation triggers rapid dysregulation and degeneration of the mouse locus coeruleus

**DOI:** 10.1101/2023.03.07.530845

**Authors:** Leslie Hassanein, Bernard Mulvey, Harris E. Blankenship, Margaret Tish, Anu Korukonda, L. Cameron Liles, Amanda L. Sharpe, Jean-Francoise Pare, Rosa M. Villalba, Xiangchuan Chen, Khan Hekmatyar, Jenny J. Yang, Daniel E. Huddleston, Seong Kang, Arielle Segal, Steven A. Sloan, Keri Martinowich, Keqiang Ye, Joseph D. Dougherty, Katharine E. McCann, Yoland Smith, Michael J. Beckstead, David Weinshenker, Alexa F. Iannitelli

## Abstract

The locus coeruleus (LC), the major source of norepinephrine (NE) in the brain, is among the first sites of pathology in both Alzheimer’s disease (AD) and Parkinson’s disease (PD), and it undergoes catastrophic degeneration later in both disorders. Dysregulation of the LC is thought to contribute to early behavioral symptoms of AD and PD such as anxiety and sleep disturbances, while frank LC loss promotes cognitive decline. However, the mechanisms responsible for this selective vulnerability are unknown. It has been suggested that neuromelanin (NM) pigment contributes to LC susceptibility, but a causal relationship has been difficult to test because rodents do not naturally produce NM. Here, we report that viral-mediated expression of human tyrosinase-induced pigmentation in male and female mouse LC neurons recapitulated key macroscopic and ultrastructural features of natural primate NM. One week of NM accumulation resulted in LC neuron hyperactivity, reduced tissue NE levels, transcriptional changes, and anxiety-like behavior. By 6 weeks, NM accumulation was associated with severe cell-autonomous LC neuron degeneration, neuroinflammation, and microglial engulfment of the pigment granules, while the anxiety-like behavior abated. These phenotypes are reminiscent of LC dysfunction and cell death in AD and PD, validating this model for studying the consequences of NM accumulation in the LC as it relates to neurodegenerative diseases.

**Significance Statement:** Alzheimer’s disease (AD) and Parkinson’s disease (PD) are the most common neurodegenerative diseases worldwide. Because therapies that cure or prevent their progression are lacking, research is focused on the identifying the earliest signs of disease as targets for diagnosis and treatment. The locus coeruleus (LC), the main source of norepinephrine (NE) in the brain, is one of the first brain regions affected in both AD and PD. Early on, LC dysregulation promotes behavioral symptoms of AD and PD, while its subsequent degeneration accelerates disease progression. Here we identify neuromelanin (NM) pigment as an LC vulnerability factor that induces neuronal hyperactivity followed by cell death. Approaches that mitigate NM accumulation and toxicity may target the earliest phases of neurodegenerative disease.

## Introduction

The neurodegenerative diseases Alzheimer’s disease (AD) and Parkinson’s disease (PD) are among the most common causes of dementia and movement disorders, respectively. The noradrenergic locus coeruleus (LC), the major source of central norepinephrine (NE), is one of the earliest brain regions to accumulate hyperphosphorylated tau (pTau) pathology in AD and α-synuclein pathology in PD (Braak et al., 2001; Del Tredici et al., 2002; Braak and Del Tredici, 2011b, a; Braak et al., 2011; Pletnikova et al., 2018; Gilvesy et al., 2022; Bueicheku et al., 2024), and it undergoes catastrophic degeneration later in both diseases (Mann et al., 1980; Bondareff et al., 1982; Iversen et al., 1983; Mann and Yates, 1983a; German et al., 1992; Zarow et al., 2003; Theofilas et al., 2017). Both clinical and animal model research has identified early LC-NE dysfunction as a trigger for early behavioral symptoms in AD and PD, such as sleep abnormalities and neuropsychiatric symptoms such as anxiety, depression, and apathy (Remy et al., 2005; Prediger et al., 2012; Ehrenberg et al., 2017; Ehrenberg et al., 2018; Sommerauer et al., 2018; Weinshenker, 2018; Butkovich et al., 2020; Gilvesy et al., 2022; Kelberman et al., 2022; Ye et al., 2022; Iannitelli et al., 2023b; Falgas et al., 2024; Korukonda et al., 2026), while subsequent frank LC degeneration predicts and exacerbates cognitive decline (Zweig et al., 1993; Heneka et al., 2006; Vazey and Aston-Jones, 2012; Rorabaugh et al., 2017; Chalermpalanupap et al., 2018; Weinshenker, 2018; Ghosh et al., 2019; Li et al., 2019; Jacobs et al., 2021; Prokopiou et al., 2022; Ye et al., 2022; Bueicheku et al., 2024).

The specific factors that make LC neurons vulnerable to pathology, dysfunction, and death in AD and PD are not fully understood, although potential contributors have been identified, including pacemaker activity, toxic NE metabolites, thin and vastly projecting unmyelinated axons, and neuromelanin (NM) (Weinshenker, 2018; Kang et al., 2020; Matchett et al., 2021; Kang et al., 2022; Iannitelli et al., 2023a; Iannitelli and Weinshenker, 2023).

NM is a dark pigment found at high abundance in catecholaminergic cells of the LC and substantia nigra pars compacta (SNc), which are also vulnerable in PD (Zecca et al., 2004; Zecca et al., 2008b; Iannitelli and Weinshenker, 2023). NM is a spontaneous byproduct of catecholamine synthesis and metabolism, and its formation likely results from an overabundance of catecholamines that cannot be sequestered into vesicles quickly enough in active neurons (Sulzer et al., 2000). In addition to catecholamine metabolites, these granules contain melanins (Bush et al., 2006), lipid droplets, protein aggregates (Sulzer et al., 2008), and heavy metals (Zecca et al., 2008b). It has been proposed that the primary function of NM is to bind and sequester these harmful compounds to mitigate potential harm to the neurons (Zecca et al., 2003). However, the buildup of NM may exacerbate neurodegeneration in AD and PD either by interfering with cellular machinery and/or through the release of previously bound toxins as it breaks down during cell death. Indeed, LC neurons containing the highest amounts of NM are disproportionately lost in PD (Mann and Yates, 1983b).

Establishing a causal relationship between NM and LC degeneration has been difficult because NM is not produced endogenously in rodents (Barden and Levine, 1983), which are the canonical animal models for studying AD and PD. While NM is thought to form non-enzymatically in human catecholamine neurons, a breakthrough came when researchers were able to drive NM formation in the SNc of mice and rats through viral-mediated expression of human tyrosinase (hTyr), the enzyme responsible for melanin production in the skin (Carballo-Carbajal et al., 2019). NM expression in the SNc resulted in neurodegeneration and subsequent motor impairments similar to those seen in other mouse models of PD. We adapted this strategy to promote NM accumulation in the mouse LC to assess the consequences of NM-mediated adaptations and neurotoxicity in a region of selective early vulnerability in AD and PD.

## Materials and Methods

### Animals

Approximately equal numbers of adult male and female mice were used for all behavioral and immunohistochemical experiments, and experimenters were blind to treatment group. Because this study was not powered to detect sex differences, and no such differences were obvious in our data, results from males and females were combined. For immunohistochemistry, behavior, and HPLC, tyrosine hydroxylase-Cre (TH-Cre) mice (B6.Cg-*7630403G23Rik^Tg(Th-cre)1Tmd^*/J, The Jackson Laboratory, # 008601) and preprodynorphin-Cre (Pdyn-Cre) mice (B6;129S-Pdyn*^tm1.1(cre)Mjkr^*/LowlJ, #027958) were used for the expression of Cre- dependent viral vectors in the LC. For translating ribosome affinity purification (TRAP) RNA-sequencing experiments, we crossed TH-Cre mice with transgenic *Slc6a2*-*eGFP*/*Rpl10a* mice (B6;FVB-Tg(*Slc6a2*-*eGFP*/*Rpl10a*)JD1538Htz/J, The Jackson Laboratory, #031151), which incorporate an EGFP/Rpl10a ribosomal fusion protein into a bacterial artificial chromosome under the *Slc6a2* (NE transporter; NET) promoter to allow for the isolation of polysomes and translating mRNAs specifically from noradrenergic neurons (Mulvey et al., 2018). Mice were group housed with sex-and age-matched conspecifics (maximum of 5 animals per cage) until one week prior to behavioral testing, and then individually housed for the subsequent week of experimentation until sacrifice. Animals were maintained on a 12:12 light:dark cycle (lights on at 0700), and food and water were available *ad libitum*, unless otherwise specified. All experiments were conducted at Emory University or the Oklahoma Medical Research Foundation in accordance with the National Institutes of Health *Guideline for the Care and Use of Laboratory Animals* and approved by the respective Institutional Animal Care and Use Committee.

### Viral vectors

To express hTyr in the LC, we developed an AAV5-EF1a-DIO-hTyr construct with assistance from the Emory Custom Cloning and Viral Vector Cores. Titers varied slightly from batch to batch but were all in the range of 10^13^ IFU/ml. For some experiments, all mice were injected with AAV5-DIO-hTyr, and TH-Cre+ animals were used as the experimental group while TH-Cre- littermates served as the controls. For other experiments, only TH-Cre+ mice were used, and mice were injected with either the AAV5-DIO-hTyr experimental virus or a comparable AAV5-DIO-EYFP control virus (Addgene, plasmid #27056).

### Stereotaxic injections

Stereotaxic infusions were performed as previously described (Tillage et al., 2020a), in a stereotaxic frame under 2.0% isoflurane anesthesia. LC infusions (unilateral or bilateral, depending on the experiment) were made at a volume of 0.5 μL/hemisphere using a 5 or 10 μL Hamilton glass syringe. The virus was infused at a rate of 0.15 μL/min, and the needle was allowed to remain in place for 5 min following the completion of each infusion. LC coordinates are AP: - 5.4mm, ML: +/- 1.2mm, and DV: -4.0mm relative to Bregma. Mice were tested at 1, 6, or 10 weeks post virus injection, depending on the experiment.

### High Performance Liquid Chromatography (HPLC)

Mice were anesthetized with isoflurane and euthanized by rapid decapitation. The pons, prefrontal cortex, and hippocampus were rapidly dissected on ice and flash-frozen in isopentane (2-Methylbutane) on dry ice. The samples were weighed and stored at -80°C until processing for HPLC by the Emory HPLC Bioanalytical Core. As previously described (Lustberg et al., 2022), tissue was thawed on ice and sonicated in 0.1 N perchloric acid (10 μl/mg tissue) for 12 s with 0.5 s pulses. Sonicated samples were centrifuged (16,100 rcf) for 30 min at 4 °C, and the supernatant was then centrifuged through 0.45 μm filters at 4000 rcf for 10 min at 4 °C. For HPLC, an ESA 5600A CoulArray detection system, equipped with an ESA Model 584 pump and an ESA 542 refrigerated autosampler was used. Separations were performed using an MD-150 × 3.2 mm C18, 3 µm column (Thermo Scientific) at 30 °C. The mobile phase consisted of 8% acetonitrile, 75 mM NaH2PO4, 1.7 mM 1-octanesulfonic acid sodium and 0.025% trimethylamine at pH 2.9. A 20 µL of sample was injected. The samples were eluted isocratically at 0.4 mL/min and detected using a 6210 electrochemical cell (ESA, Bedford, MA) equipped with 5020 guard cell. Guard cell potential was set at 475 mV, while analytical cell potentials were −175, 100, 350 and 425 mV. The analytes were identified by the matching criteria of retention time measures to known standards (Sigma Chemical Co., St. Louis MO). Compounds were quantified by comparing peak areas to those of standards on the dominant sensor.

### Immunohistochemistry

Mice were euthanized with an overdose of isoflurane and were transcardially perfused with cold 4% PFA in 0.01 M PBS for light microscopy and 4% PFA + 0.1% glutaraldehyde for electron microscopy. After extraction, brains were post-fixed overnight in 4% PFA at 4°C and then transferred to a 30% sucrose/PBS solution for a minimum of 72 h at 4°C. Brains were embedded in OCT medium (Tissue-Tek) and sectioned by cryostat into 40-μm-thick coronal sections at the level of the LC, anterior cingulate cortex (ACC), and hippocampus. Sections were blocked for 1 h in 5% normal goat serum (NGS) in 0.01 M PBS/0.1% Triton-X permeabilization buffer and then incubated for 24 h at 4°C in NGS blocking buffer with primary antibodies listed in **Supplemental Table 1**. Following washes in 0.01 M PBS, sections were incubated for 1.5 h in blocking buffer including Nissl and secondary antibodies listed in **Supplemental Table 1**. After washing, sections were mounted onto Superfrost Plus slides and cover-slipped with Fluoromount-G plus DAPI (Southern Biotech, Birmingham, AL).

### Fluorescence Microscopy

For the catecholaminergic marker tyrosine hydroxylase (TH), astrocyte marker glial fibrillary acidic protein (GFAP), ionized calcium-binding adaptor molecule 1 (Iba1) and LC terminal marker NE transporter (NET), listed in **Supplemental Table 1**, immunofluorescent images were acquired as z-stack images (10 z-stacks; pitch: 0.1 µm) at 20x magnification with uniform exposure parameters and full-focused on a Keyence BZ-X800 microscope system. Brightfield images of NM pigment granules were also obtained on the Keyence BZ-X800 microscope at 20x magnification. Following convention, these images are oriented with the dorsal direction up and the ventral direction down.

### Cell counts

Animals were perfused as described in the immunohistochemistry section above. Brains were sectioned at 40 μm on a cryostat, and sections were processed for immunohistochemistry to stain for TH and Nissl. Brightfield images to capture NM expression were transformed into fluorescence images using hDAB deconvolution in QuPath v.0.7.0 (Bankhead et al., 2017) and fused with fluorescence images of TH and Nissl. Cell counts were quantified manually using FIJI/ ImageJ image overlays. Cells were defined using either Nissl or TH and then counted within the region of interest delineating the LC. LC borders were defined using TH and NM. Me5 cells, which fluoresce brightly in all channels and are much larger and rounder than LC cells, were excluded from the boundaries of analysis. Parameters used to define cells were set for each animal and used consistently across all sections for that animal.

### Inflammatory marker quantification

One representative, atlas-matched image per channel (GFAP, Iba1) was selected based on anatomical landmarks and TH labeling in the LC. To quantify inflammatory markers and NET in the hippocampus and prelimbic cortex, one atlas-matched image per animal was selected based on anatomical structure. Quantification of inflammatory markers was performed using FIJI/ImageJ (NIH). All images were processed using an identical analysis pipeline to ensure consistency across samples. Images were converted to 8-bit grayscale and background subtracted using a rolling ball radius of 25 pixels with the sliding paraboloid option enabled. Brightness and contrast were standardized across all images prior to thresholding. A fixed threshold was applied to all images within a given analysis set, determined empirically to best capture signal while minimizing non-specific labeling and background. Signal quantification was then performed using the “analyze particles” function with a minimum particle size of 5 pixels to exclude noise, and the “limit to threshold” option enabled to restrict analysis to signal-positive pixels. Measurements include area and integrated density (IntDen). A single summary value was extracted representing the total integrated density and total area of threshold signal across the image.

### Electron microscopy

#### Tissue collection and processing

All brains prepared for electron microscopy were perfused with a Ringer’s solution and a mixture of paraformaldehyde (4%) and glutaraldehyde (0.1%), before being post-fixed for 14 h in 4% paraformaldehyde and cut in 60 µm-thick coronal sections with a vibrating microtome. Selected sections that included the LC were immunostained with specific antibodies against TH (dilution 1:2000; Millipore, AB152; RRID: AB_390204), IBA-1 (dilution 1:500; Abcam; Ab5076; RRID: AB_2224402) and GFAP (dilution 1:5000; Abcam AB7260; RRID: RRID:AB_305808) according to the avidin biotin complex method (ABC-Vectastain Standard kit, Vector Labs), using DAB as chromogen for the peroxidase reaction. Following the DAB reaction, sections were transferred to a phosphate buffer solution (PB, 0.1 M, pH 7.4).

#### Transmission electron microscopy (TEM)

The tissue processed for TEM was postfixed in a 1% osmium tetroxide solution for 20 min. Following washes in PB, and then the samples were dehydrated in alcohol solutions (50–100%) before being placed in propylene oxide. Uranyl acetate (1%) was added to the 70% alcohol to increase contrast in the electron microscope. The dehydrated sections were embedded in epoxy resin (Durcupan, ACM; Fluka, Buchs, Switzerland) for 12 h, mounted onto oil-coated slides and cover-slipped before being baked at 60°C for 48 h. Blocks of LC tissue were then taken out from the slide, glued on resin blocks and cut in ultrathin sections (70 nm-thickness) using an ultramicrotome (Ultra-cut T2; Leica, Germany) and mounted onto single slot Pioloform-coated copper grids. Grids were then examined with a transmission electron microscope (EM; Jeol; Model 1011) coupled with a CCD camera (Gatan; Model 785) controlled with DigitalMicrograph software (Gatan; version 3.11.1).

For ultrastructual comparisons of hTyr-induced NM in mice and naturally produced NM in primates, three brainstem sections at the level of LC from a 16-year-old rhesus macaque (from the monkey brain tissue bank in the Y. Smith lab) were processed according to the same procedure and examined in the electron microscope.

#### Tissue processing and image acquisition for array tomography/scanning electron microscopy (AT/SEM)

Samples of GFAP- and IBA-1-immunostained LC were used for 3D EM analysis. After stopping the DAB reaction with PBS washes, circular punches of tissue (1-mm diameter) were taken from the LC immunostained sections and shipped to the Oregon Health Science Center University Microscopy Core (Portland, Oregon) in 4% paraformaldehyde for AT/SEM processing. Samples were first rinsed in 0.1M sodium cacodylate buffer and stained with tannic acid (1%) in 0.1 M sodium cacodylate for 15 min. Tissue was then incubated in osmium tetroxide (2%) reduced with potassium ferricyanide (1.5%) in 0.1 M sodium cacodylate for 1 hr, followed by overnight incubation in uranyl acetate (1%) at 4 °C. Then, samples were treated with lead aspartate at 60 °C for 1 hr, dehydrated through acetone series (50%, 75% 85%, 95%, 100%), and infiltrated with resin overnight. Finally, samples were further infiltrated with resin and polymerized in the oven at 60℃. Ultrathin sections (70 nm) were cut using a Leica Artos 3D ultramicrotome equipped with a Leica AT-4 diamond knife and collected as ribbons onto silicon chips. The chips were mounted on a stub and imaged using a Helios 5 UC scanning electron microscope (Thermo Fisher Scientific) with Maps™ software array tomography module.

Regions of interest were acquired at 4-nm pixel resolution (10,240 x 10,240 pixels) using the CBS detector. Image stacks (∼150 to 200 serial images per stack) were aligned in FIJI using the *Linear Stack Alignment with SIFT* plugin, with the expected transformation set to affine. The aligned images were then imported into Amira (Thermo Fisher Scientific), to extract a 6144 x 6144 pixels subvolume. They were then imported into the *Reconstruct* (NIH) software (https://synapseweb.clm.utexas.edu/software-0), the section thickness (65-70nm) and the pixel size (0.0045μm) were adjusted before the segmentation, and the contours of the different elements analyzed were manually traced with different color code lines in each serial electron micrograph.

### Neuromelanin-magnetic resonance imaging (NM-MRI)

Mice described above were anesthetized, monitored, and scanned using a Bruker Biospec 7 T MRI scanner. Before commencing the scanning, animals were placed in an induction chamber connected to an isoflurane anesthesia unit. Following anesthesia induction with isoflurane (3-5% in oxygen), mice were transferred to the MRI cradle and placed on an imaging platform (warmed to 37° C) where anesthesia was maintained with a nosecone (1-3% isoflurane) utilizing the SomnoSuite (Kent Scientific, Somnosuite, CT) small animal anesthesia delivery system. Upon induction of anesthesia, the animal’s eyes were lubricated with a sterile ophthalmic ointment. A thermocouple rectal probe was placed in the rectum to monitor core temperature. Respiratory rate was also monitored (SAI instruments Inc, NY), with a target rate of 50-65 breaths/min. Light paper tapes were used to minimize the movement of the animal body in addition to the stereotactic setup.

MRI scanning was performed for ∼90 min using a Biospec Bruker 7T/21 cm MRI (Bruker Billerica, Manning Park Billerica, MA) inner diameter horizontal bore magnet using a specialized large 86 mm *^1^H transmit-receive volume coil* radio frequency coil with inner diameter of 72 mm coupled with 4 channel mouse array brain coil.(^1^H receive-only 2 x 2 mouse brain surface array coil). All localizer and preparatory scans were performed before the NM-MRI scans. A baseline localizer scan was obtained along the three orthogonal directions using the gradient echo scan (Gradient Recalled Echo). Then, a B_0_ map was obtained for the heart to correct any inhomogeneities. Three additional reference scans are acquired and served as a reference to setup the appropriate slices for NM-MRI.

NM-MRI images were acquired using a magnetization transfer (MT) prepared gradient echo pulse sequence with echo time = 3.654 ms, repetition time = 31.0 ms, averages = 10, voxel size = 0.125 mm x 0.125 mm x 0.250 mm, an MT preparation pulse with 14.6 ms duration, pulse amplitude 8.21 μT, and MT flip angle 768.0°, and an acquisition time 1 h 33 min 55 s.

### Translating Ribosome Affinity Purification (TRAP)

To obtain adequate quantities of RNA for sequencing, samples from two 6-8 month-old, same-sex and treatment TH-Cre+, *Slc6a2*-*eGFP*/*Rpl10a+* mice were pooled to form a biological replicate by dissecting out the hindbrain posterior to the pontine/hypothalamic junction (cerebellum was discarded). Six biological replicates were collected per treatment group. Each replicate was homogenized and TRAP was performed as described (Mulvey et al., 2018; Iannitelli et al., 2022), resulting in LC-enriched “TRAP” samples and whole-hindbrain “input” samples. RNA was extracted using Zymo RNA Clean & Concentrator-5 kit, and subsequently sent for polyA-enriched library preparation and Illumina sequencing by NovoGene to a minimum depth of 20 million fragments per sample. Forward and reverse sequencing files from each replicate were aligned to the mouse genome (mm10) using STAR alignment, and counts were obtained using FeatureCounts in R Bioconductor. Two samples from saline-treated mice were removed from analysis because the TRAP protocol failed to enrich *Slc6a2* above a 10-fold change, a quality control threshold observed in all other saline-treated samples. Sequencing data are available on NCBI GEO (GSE226827).

#### DE analysis

Gene counts data were imported into R 4.4.1 on a Macintosh for analysis using *edgeR* and the *voomWithQualityWeights* (Mulvey et al., 2023) implementation as incorporated into *voomLmFit* (Chen et al., 2025). Counts data were imported and distributions of counts visualized to determine a low-count cutoff, which we set to 70, requiring at least 5 samples to exceed this threshold for a given gene. A total of 13,213 genes were therefore retained for downstream differential expression analyses. TMM normalization factors were then calculated to account for sequencing library size (Mulvey et al., 2023). DE was performed using a model covering 4 groups: EYFP input, EYFP TRAP (LC), hTyr input, and hTyr TRAP, allowing us to compare both TRAP vs. input (within condition) TRAP vs. TRAP (across conditions) using the same fitted data. Our model was set up as *∼0+group* and initialized in *voomLmFit* with sample weights enabled (as is implicit in the predecessor, *voomWithQualityWeights*). *voomLmFit* acts as a wrapper around multiple DE pipeline steps in that it executes *duplicateCorrelation* (Smyth et al., 2005) at the level of replicate to correct for multiple samples having been collected from the same mice (every given homogenate was split into an input and a TRAP fraction), and further executes *voomWithQualityWeights* a second time to refine parameter estimates and *lmFit* to fit the model. We subsequently used *topTable* to collect full DE stats for each of the four comparisons of interest.

#### Gene Set Enrichment Analysis (GSEA)

To test for enrichment of DE genes in ontology and literature-derived sets, we utilized the threshold-free GSEA method (Subramanian et al., 2005) and its accompanying gene set catalogs (curated/C2, regulatory targets/C3, ontologies/C5, oncogenic/C6, and cell type signatures/C8) in mSigDB version 2024.1 mouse, implemented using the R package *fGSEA* (Korotkevich et al., 2021). We additionally collated several additional TF-target gene sets to test for enrichment: ChEA 2022 resource (Keenan et al., 2019), TRRUST (Han et al., 2018), those mined from publication supplements in Rummagene (Clarke et al., 2024), and as curated in the Enrichr (Xie et al., 2021) package (Gene Expression Omnibus-mined results of TF perturbation experiments and TF protein-protein interactions). We input the signed DE *t* statistics for use as the DE gene ranking values to *fGSEA* with analysis parameters as follows: gene set sizes 15-500; *eps* (specifying range over which to calculate p-values) set to 0 to return a p-value for all tests; and 50,000 permutations for initial p-value estimations. For mSigDB sets, *fGSEA*’s *collapsePathways* function was then used to consolidate results to represent the strongest/most specific enrichments from overlapping sets (in other words, the most granular set based on *de novo* calculation of a hierarchy of sets).

For analysis of DEG data reported in (Mathys et al., 2019), we utilized the provided DE analysis results from Supplemental Table 2 for excitatory neurons, with the signed ‘MixedModel.z’ statistic used as the DE ranking variable. We utilized the same collated TF-target gene sets as used for GSEA analysis of our data. fGSEA parameters were gene set sizes 15-500; *eps* set to 0 to return a p-value for all tests; and 10,000 permutations for initial p-value estimations. While we performed analyses using the full corpus of TF-target sets to ensure comparable multiple testing correction, we only examined TF-target sets corresponding to *ATF6* and *XBP1* for interpretation of this analysis.

#### STRING Protein-Protein Interactors for Xbp1 and Atf6

To determine physical interactors of *Xbp1* and *Atf6*, we queried mouse protein-protein interactions indexed in STRINGdb v12.0 by searching the two mouse gene symbols. Subsequently, we filtered to interactions listed as “high confidence (>0.7)” and downloaded the list of 82 mouse protein interactors meeting this criterion, 55 of which were included in the DE analysis. We then examined the hTyr-LC vs. EYFP-LC DE table for significant DEGs among these interactors. The result of this STRINGdb query is included in **Supplemental Table 4**.

### Electrophysiology

Mice were deeply anesthetized with isoflurane and decapitated. Brains were rapidly removed and sectioned in ice-cold cutting solution containing (in mM): 110 choline chloride, 2.5 KCl, 1.25 Na_2_PO_4_, 0.5 CaCl_2_, 10 MgSO_4_, 25 glucose, 11.6 Na-ascorbate, 3.1 Na-pyruvate, 26 NaHCO_3_, 12 N-acetyl-L-cysteine, and 2 kynurenic acid. Horizontal (220 µm) slices containing the LC at the level of the mesencephalic trigeminal tract neurons (ME5) were collected and transferred to a holding chamber containing artificial cerebrospinal fluid (aCSF) containing (in mM) 126 NaCl, 2.5 KCl, 1.2 MgCl_2_, 2.4 CaCl_2_, 1.2 NaH_2_PO_4_, 21.4 NaHCO_3_, and 11.1 glucose plus 1 Na ascorbate, 1 Na pyruvate, 6 N-acetyl-L-cysteine and MK-801 to minimize excitoxic effects induced by sectioning. Slices recovered for 30 min at 32°C followed by at least 30 min at room temperature prior to recording.

Slices were transferred to a recording chamber where they were perfused with warmed aCSF (inline heater, 32-34°C, Warner Instruments) at a rate of approximately 2 mL/min via gravity or a peristaltic pump (Warner Instruments). Slices were visualized under Dödt gradient contrast (DGC) optics on an upright microscope (Nikon FN1 or Zeiss Examiner D1). Putative LC-NE neurons were identified first by location (immediately medial to ME5 neurons, rostral to the fourth ventricle) and large (>20 µm) cell bodies, followed by visual presence of NM or EYFP fluorescence (controls). Electrophysiological parameters for LC-NE neuron identification included slow, irregular spontaneous action potential generation, presence of A-type potassium currents and a rebound delay after a hyperpolarizing step (-100 pA) in current clamp and low input resistance (approximately 100 MΩ, -60 V_hold_) (Williams et al., 1984). Thin wall glass (World Precision Instruments or Warner Instruments) was used for cell-attached and whole-cell recordings, and displayed resistances of 2.5-3 MΩ when filled with an intracellular solution containing (in mM): 135 K gluconate, 10 HEPES, 5 KCl, 5 MgCl_2_, 0.1 EGTA, 0.075 CaCl_2_, 2 ATP, 0.4 GTP, pH 7.35. The liquid junction potential was calculated to be -17 mV, and was not corrected. Cell attached recordings were conducted using a Na-HEPES based intracellular solution (plus 20 mM NaCl, 290 mOsm/L, pH 7.4) (Branch and Beckstead, 2012). Frequency-current (F-I) relationships were calculated as spikes per second across the 2 second depolarization step. Gain was calculated as the slope of the steady state F-I curve, from 0-250pA. Cell-attached frequency was calculated as spikes per second during a 30 second sweep.

### Behavioral assays

Behavioral assays were performed in the following order, from least to most stressful.

#### Sleep latency

Latency to sleep was measured as the duration of time it took for the mouse to fall asleep following mild stimulation (gentle handling). A sleep bout was defined as 2 min of uninterrupted sleep behavior, followed by 75% of the following 8 min (Hunsley and Palmiter, 2004). This assay has been previously validated using EEG recordings (Porter-Stransky et al., 2019). Sleep testing began at 9:00 A.M., 2 h into the light cycle when mice typically engage in sleep behavior naturally. Videos were recorded for each session and scored by a blinded observer. Previous research by our group and other has revealed that increasing LC-NE transmission decreases sleep latency, while NE depletion has the opposite effect (Porter-Stransky et al., 2019).

#### Novelty-suppressed feeding

Chow was removed from individual home cages 24 h prior to behavioral testing, which was conducted in the early afternoon. Mice were moved to the test room under red light and allowed to habituate for 2 h prior to the start of the test. Individual mice were placed in a novel arena (10” × 18” × 10”) with a single pellet of standard mouse chow located in the center. The latency to feed, operationally defined as grasping and biting the food pellet, was recorded using a stopwatch. Mice that did not feed within the 15-min period were assigned a latency score of 900 s (Tillage et al., 2020b). We have shown that increasing NE promotes anxiety-like behavior in this task reflected by longer latencies to eat in the novel environment, while decreasing attenuates anxiety and reduces latency to eat (Lustberg et al., 2020).

#### Fear conditioning and context testing

Fear-conditioning training and subsequent contextual fear testing is a widely used assessment of associative memory, particularly for an environment in which an aversive stimulus (footshock) was previously administered. This method has been described previously by our group and others and is sensitive to changes in NE (Murchison et al., 2004; Chalermpalanupap et al., 2018; Butkovich et al., 2020). All conditioning and testing were conducted in the early afternoon. Mice were placed in a fear-conditioning apparatus (7 in. x 7 in. x 12 in.; Coulbourn Instruments) with a metal shock grid floor. Following 3 min of habituation, three conditioned stimulus (CS)-unconditioned stimulus (US) pairings were presented with a 1-minute intertrial interval. The CS was a 20-second, 85 dB tone which co-terminated with the US, a 2-s, 0.5 mA footshock (Precision Animal Shocker, Colbour Instruments). The following day, the context test was conducted by placing animals back into the fear conditioning chamber without the administration of CS-US pairings. Freezing behavior was measured as a readout of memory for the fear-associated context.

### Statistical analyses

Cell counting, immunofluorescence, and HPLC measurements of catecholamine concentrations were compared between hTyr-injected and control groups using unpaired t-tests in GraphPad Prism. Similarly, behavioral assessment relied on unpaired t-test comparison between groups for latency to feed in the novelty-suppressed feeding task, latency to fall asleep, and freezing in the contextual fear assay. Welch’s t-tests were used for comparing groups with unequal variances. Statistical analyses of RNA sequencing data are described above.

## Results

### Viral infusion of hTyr induces NM-like pigmentation in the LC of mice

To recapitulate endogenous NM found in the human LC, we adapted a viral vector-mediated approach to drive neuronal pigmentation in mice. TH-Cre+ and TH-Cre- mice were stereotaxically infused bilaterally with AAV5-DIO-hTyr in the LC to drive viral expression of hTyr (**Fig. 1a**), which is known to produce pigment when introduced in rodent SNc DA neurons (Carballo-Carbajal et al., 2019). We found that hTyr expression likewise drove pigmentation of the LC in Cre+ (**Fig. 1b**) but not in Cre- control mice (**Fig. 1c, d**) as early as 1-week post-infusion. NM-MRI revealed a hyperintense signal overlying the location of the LC only on the hTyr side of mice that received hTyr virus in one hemisphere and AAV5-DIO-EYFP control virus in the other hemisphere (**Fig. 1e**). Additionally, we confirmed the presence of melanin in this pigment using Fontana-Masson staining (**Fig. 1f**). We then utilized electron microscopy to compare the pigment granules in our rodent model (**Fig. 1g**) with endogenous NM granules in tissue obtained from a 16-year-old Rhesus macaque (**Fig. 1h**). Ultrastructural inspection revealed that many of the components known to comprise endogenous NM, including tightly packed melanin pigments and lipid droplets, were also present in granules from our rodent model. Together, these results validate the use of viral-mediated hTyr expression to induce NM-like pigmentation in the mouse LC.

**Fig. 1.**
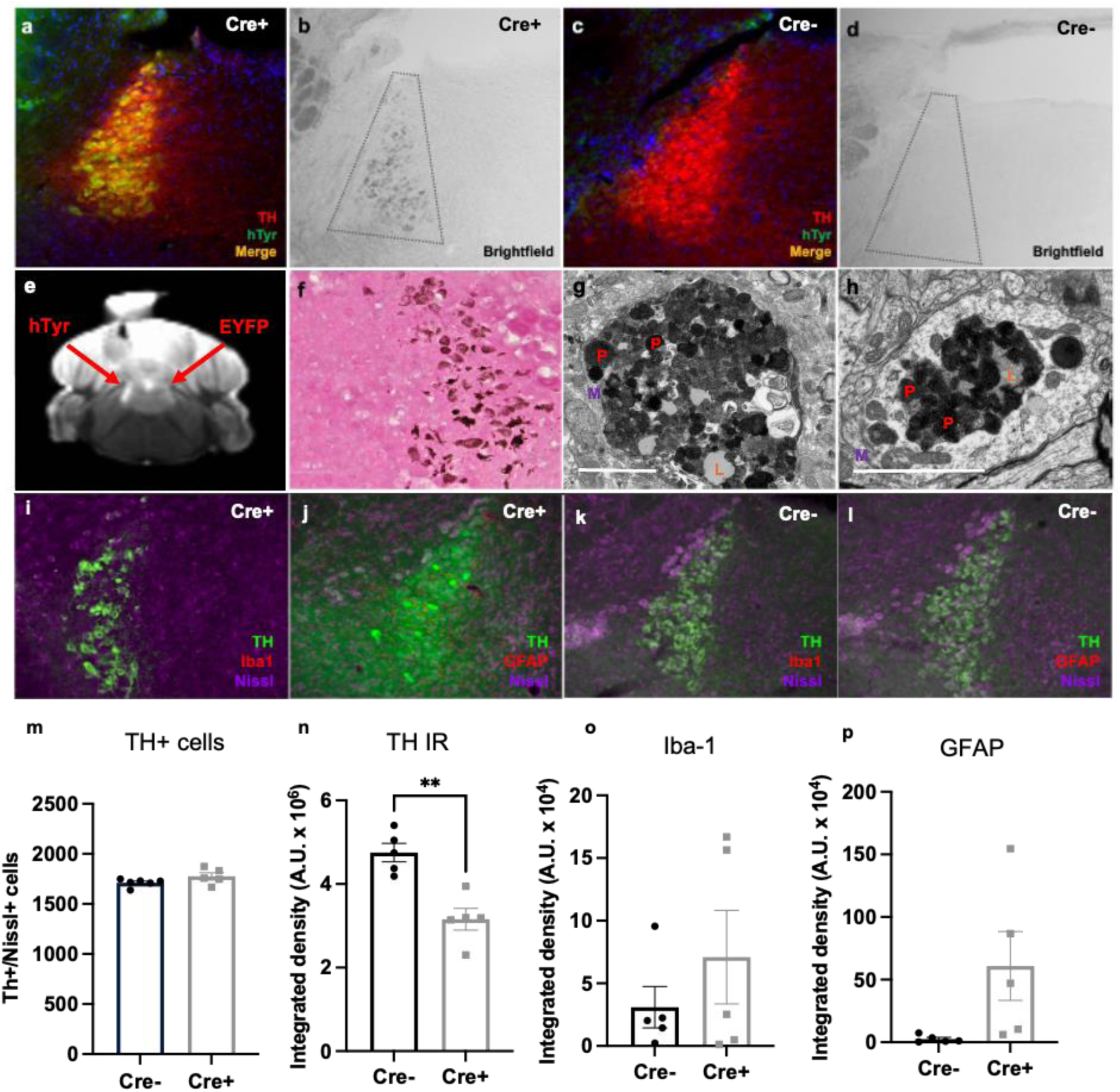
hTyr expression drives NM accumulation in the LC at 1 week. TH-Cre- or TH-Cre+ mice received bilateral LC infusion of AAV5-DIO-hTyr and were assessed for NM accumulation and LC neuron integrity 1 week later. Shown are representative images of TH and hTyr immunoreactivity (**a**, **c**), NM in brightfield (**b**, **d**), NM-MRI contrast (**e**), Fontana-Masson staining, a NM granule from the LC of a TH-Cre+ mouse expressing hTyr in the LC (**g**), a NM granule from the LC of a 16-year old rhesus macaque (**h**), TH, Iba-1, and Nissl (**i**, **j**), and TH, GFAP, and Nissl (**k**, **l**). Quantification (mean ± SEM) of TH+/Nissl+ cells (**m**), total TH immunoreactivity (**n**), Iba-1 immunoreactivity (**o**), and GFAP (**p**) immunoreactivity are shown below (N=5-6/group). p, pigment (red); l, lipid droplet (orange), scale bar = 5 μM. Immunofluorescence, brightfield, and Fontana-Masson images were acquired at 20X. **p<0.01.

### NM results in reduced TH immunoreactivity, but not TH+ cell bodies in the LC at 1 week with no changes in inflammatory markers

To assess whether pigmentation impacted cell body integrity at 1 week post-injection, we performed a count of TH+ and Nissl+ cells in the LC and found no difference between TH-Cre+ and TH-Cre- mice that received hTyr virus (t_(9)_ = 1.73, p = 0.11) (**Fig. 1i-m**). However, despite a handful of strongly fluorescent TH+ cells in the LC of TH-Cre+ mice, overall TH immunoreactivity in the region was significantly reduced (t_(8)_ = 4.66, p = 0.002) (**Fig. 1i, j, n**), consistent with the NM-induced loss of TH observed in the SNc of rats expressing hTyr (Carballo-Carbajal et al., 2019). Although both microglial (Iba-1) and astroglial (GFAP) inflammation tended to be higher in the LC region of NM-expressing mice as compared to controls, there was a high degree of individual variability and the group differences were not significant Iba-1: t_(8)_ = 0.98, p = 0.36; GFAP: t_(4)_ = 2.12, p = 0.10) (**Fig. 1o, p**).

### Effects of NM on noradrenergic fiber integrity, neuroinflammation, and catecholamine levels in LC projection fields at 1 week

Because LC axon/terminal degeneration precedes cell body loss in clinical AD and PD, as well as rodent models of these diseases (Chalermpalanupap et al., 2013; Rorabaugh et al., 2017; Butkovich et al., 2020; Doppler et al., 2021a; Gilvesy et al., 2022), we also assessed NET immunoreactivity, a marker of LC fiber integrity, in two major forebrain projection regions of the LC (hippocampus and prelimbic cortex). There was a borderline significant reduction in NET immunoreactivity in the hippocampus (t_(10)_ = 2.12, p = 0.06) (**Fig. 2a, d, m**) but not prelimbic cortex (t_(10)_ = 0.24, p = 0.82) (**Fig. 2g, j, p**) of TH-Cre+ mice compared to TH-Cre- controls. This selective vulnerability of hippocampal-projecting fibers compared to cortical-projecting fibers is reminiscent of the pattern we observed in TgF344-AD rats that accumulate early pTau in the LC (Rorabaugh et al., 2017). Next, we assessed neuroinflammatory markers in LC projection regions and found significant elevation of Iba-1 in the hippocampus (t_(10)_ = 3.08, p = 0.01) (**Fig. 2b, e, n**) and prelimbic cortex (t_(5)_ = 4.29, p = 0.007) (**Fig. 2h, k, q**), with no changes in GFAP (hippocampus: t_(10)_ = 1.29, p = 0.23, **Fig. 2e, f, o**; prelimbic cortex: t_(5)_ = 1.10, p = 0.32, **Fig. 2i, l, r**), indicating an increase in microglial, but not astrocyte, reactivity.

**Fig. 2.**
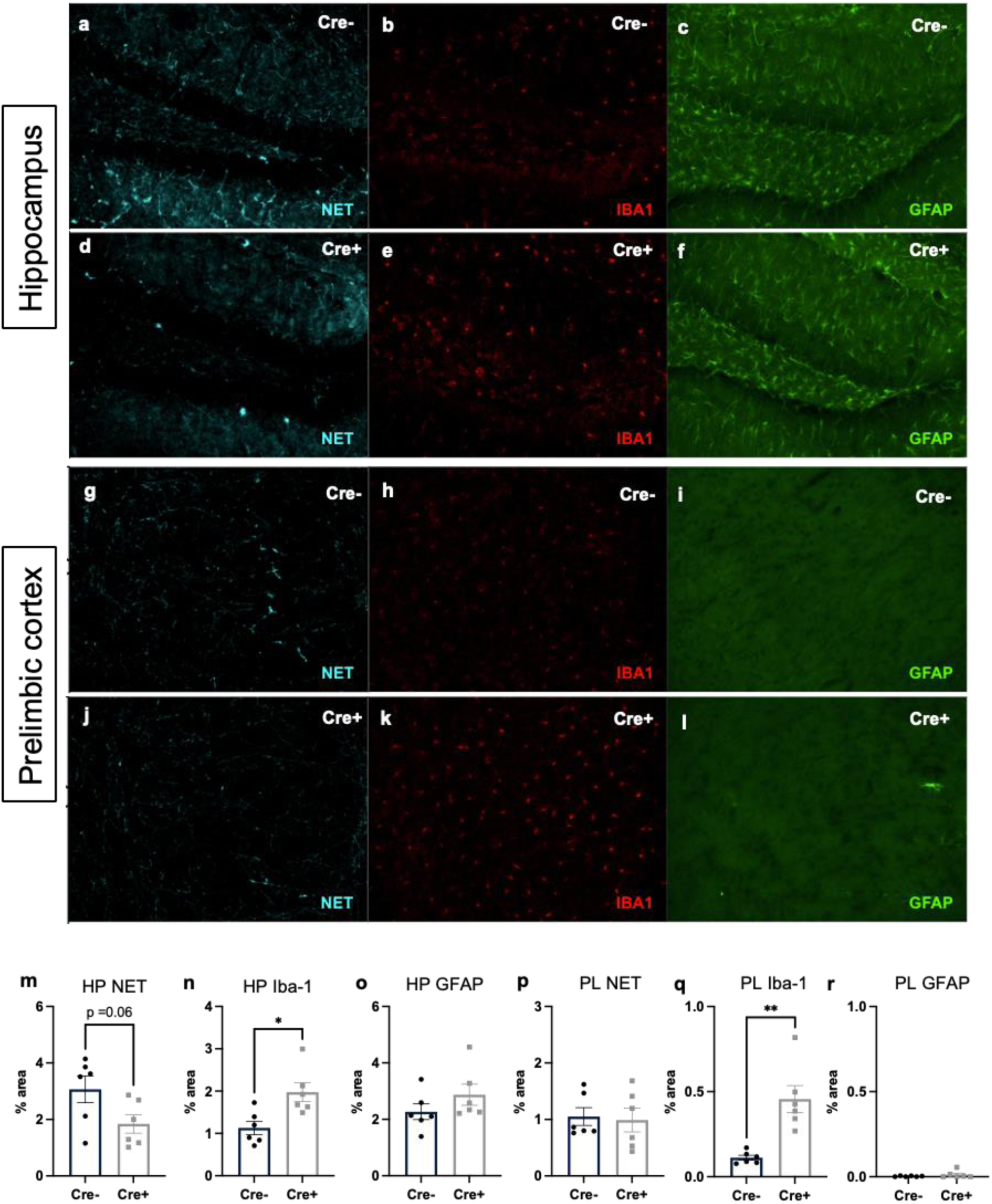
NM accumulation in the LC is associated with changes in noradrenergic innervation and neuroinflammation in the hippocampus and prelimbic cortex. TH-Cre- or TH-Cre+ mice received bilateral LC infusion of AAV5-DIO-hTyr and were assessed for NET, Iba-1, and GFAP immunoreactivity 1 week later. Shown are representative images of NET (**a**, **d**), Iba-1 (**b**, **e**), and GFAP (**c**, **f**) in the hippocampus and in in the prelimbic cortex (**g**-**l**). Data (mean ± SEM) are quantified below (**m**-**r**) (N=6/group). HP, hippocampus; PL, prelimbic cortex. Immunofluorescence images were acquired at 20X. *p<0.05, **p<0.01.

Given the loss of LC axons/terminals in the hippocampus and the reduction of TH immunoreactivity, we assessed the impact of hTyr-driven pigmentation on tissue levels of NE and its primary metabolite 3-Methoxy-4-hydroxyphenylglyucol (MHPG) in a separate group of TH-Cre+ mice that received either bilateral hTyr virus or EYFP control virus. NE was dramatically reduced in the pons, where LC cell bodies reside (t_(10)_ = 10.14, p < 0.0001) (**Fig. 3a**), as well as in the hippocampus (t_(10)_ = 9.735, p < 0.0001) (**Fig. 3d**) and prefrontal cortex (PFC; t_(10)_ = 7.607, p < 0.0001) (**Fig. 3g**) in the hTyr mice. Levels of the primary NE metabolite MHPG were also reduced in the pons (t_(10)_ = 7.53, p < 0.0001) (**Fig. 3b**), hippocampus (t_(10)_ = 3.67, p = 0.004) (**Fig. 3e**), and PFC (t_(10)_ = 5.15, p = 0.0004) (**Fig. 3h**). Because NE depletion was more severe than MHPG in LC terminal regions, the MHPG:NE ratio, which provides an estimate of NE turnover, was significantly increased in the hippocampus (t_(10)_ = 4.39, p = 0.001) (**Fig. 3f**) and PFC (t_(10)_ = 2.91, p = 0.016) (**Fig. 3i**) of NM-expressing mice compared to controls. Levels of other monoamine neuromodulators, including dopamine (DA), serotonin (5-HT), and their respective metabolites, were unchanged with the exception of an increase in DA turnover (DOPAC:DA ratio) in both the pons (t_(10)_ = 3.288, p = 0.009) and PFC (t_(10)_ = 2.905, p = 0.016), suggesting that dysfunction resulting from pigmentation of the LC was largely confined to the noradrenergic system at 1 week post-injection (**Supplemental Table 2**). Because LC neurons produce and release DA as well as NE (Ranjbar-Slamloo and Fazlali, 2019), it is possible that the increased DA turnover can also be traced to LC hyperactivity.

**Fig. 3.**
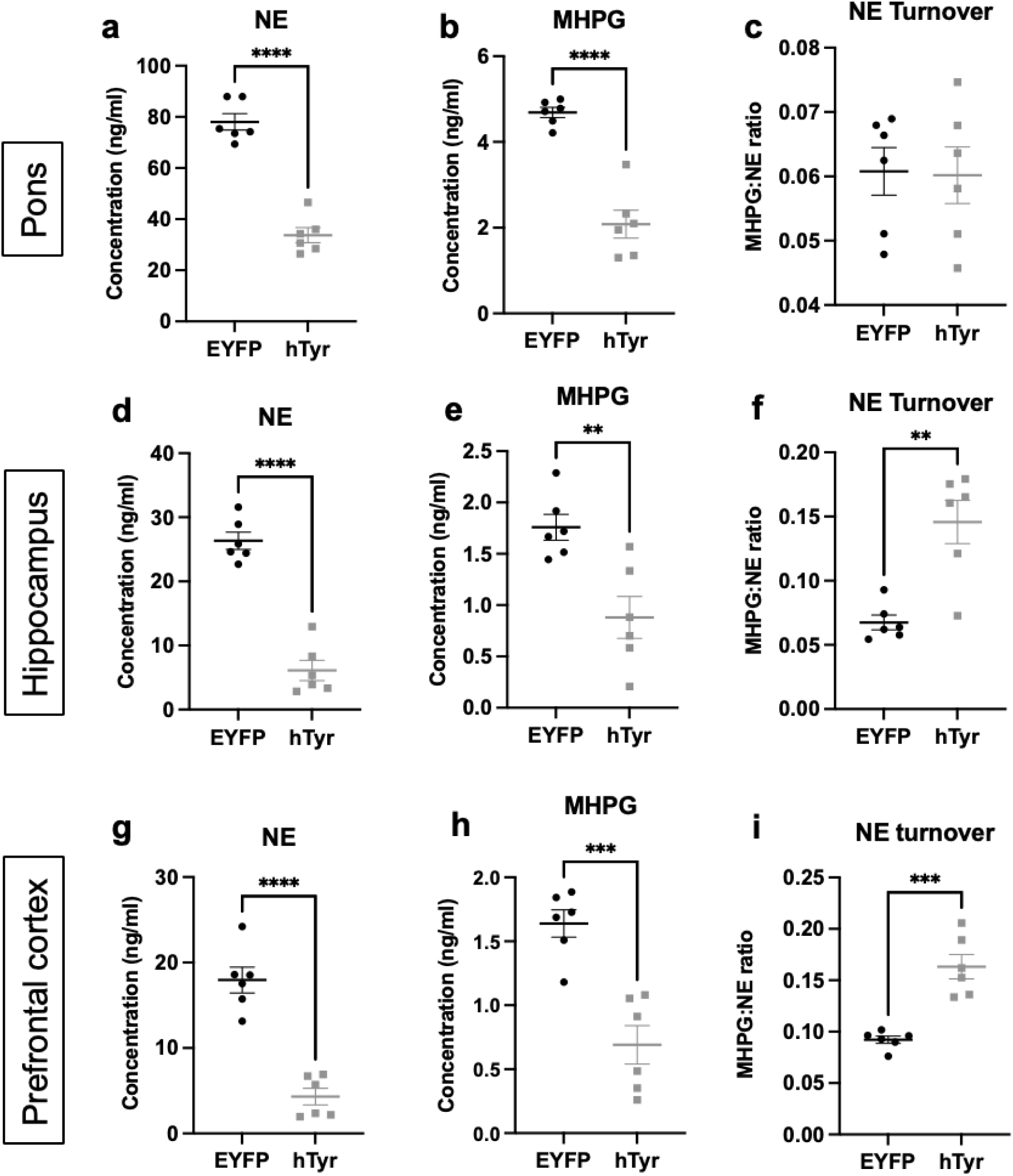
hTyr-induced NM decreases tissue NE and MHPG levels and increases turnover at 1 week. TH-Cre mice received bilateral tereotaxic infusion of AAV-DIO-hTyr or AAV-DIO-EYFP into the LC, and tissue NE levels, MHPG levels, and NE turnover (MHPG:NE ratio) were measured 1 week later by HPLC. Shown is (mean ± SEM) in the pons (**a**-**c**), hippocampus (**d**-**f**), and prefrontal cortex (**g**-**i**). N=6 per group. **p<0.01, ***p<0.001, ****p<0.0001.

### NM-positive LC neurons are hyperactive at 1 week

Given the increase in NE turnover, we speculated that at 1 week post-viral infusion, LC neurons burdened with NM were hyperactive. Using *ex vivo* slice electrophysiology, we found that both spontaneous pacemaker (**Fig. 4a**) and current-evoked (**Fig. 4b**) firing were significantly elevated in LC neurons that contained NM from TH-Cre+ mice that received bilateral hTyr virus compared to those from mice that received EYFP virus or other controls (those that received hTyr virus but did not display visible pigment, no virus). For spontaneous firing, one-way ANOVA showed a significant difference between groups (F_2,149_ = 24.02, p < 0.0001), and post hoc analysis revealed that the NM-containing LC neurons were hyperactive compared to EYFP controls (q = 8.81, p< 0.0001) and other controls (q = 9.26, p < 0.0001). For evoked firing, two-way ANOVA showed a main effect of treatment (F_1,99_ = 25.35, p < 0.0001), current injection (F_14,1386_ = 236.20, p < 0.0001), and a treatment x current injection interaction (F_14,1386_ = 5.49, p < 0.0001). Post hoc analysis revealed that the NM-containing LC neurons were hyperactive compared to EYFP controls at 50 pA (t = 3.40, p < 0.01), 100 pA (t = 4.19, p < 0.001), 140 pA (t = 4.74, p < 0.0001), 180 pA (t = 5.46, p < 0.0001), 220 pA (t = 5.27, p < 0.0001), 260 pA (t = 5.69, p < 0.0001), 300 pA (t = 4.86, p < 0.0001), 340 pA (t = 4.59, p < 0.0001), 380 pA (t = 3.82, p < 0.01), and 500 pA (t = 2.69, p < 0.05).

**Fig. 4.**
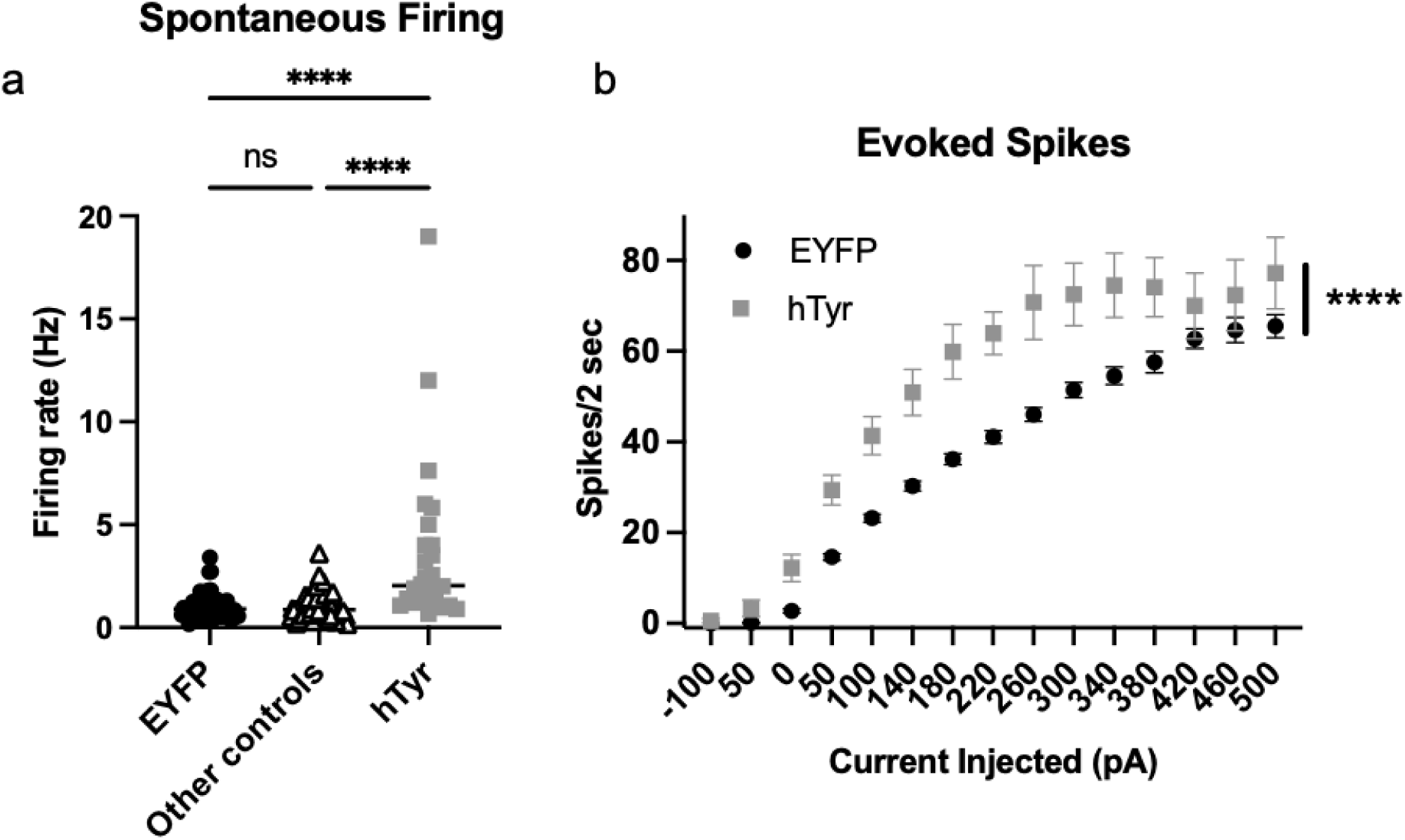
hTyr-induced NM accumulation causes LC hyperactivity at 1 week. TH-Cre mice received bilateral stereotaxic infusion of AAV-DIO-hTyr or AAV-DIO-EYFP into the LC, and LC neuron activity was assessed 1 week later by slice electrophysiology. Shown is (**a**) spontaneous firing (hTyr, N=26; EYFP, N=55; other controls including non-NM expressing neurons in the hTyr group, N=71) and (**b**) (mean ± SEM) current injection-evoked spikes (hTyr, N=24; EYFP, N=77). ****p<0.0001.

### The presence of NM in the LC drives novelty-induced anxiety behavior at 1 week

Several early behavioral symptoms of AD and PD are regulated by the LC-NE system, and thus are sensitive to the early LC dysfunction that precedes neurodegeneration (Weinshenker, 2018). In particular, we have shown that LC hyperactivity and/or NE signaling is associated with increased anxiety-like behavior in rodent models of AD and PD (Butkovich et al., 2020; Kelberman et al., 2022; Iannitelli et al., 2023b; Kelberman et al., 2023; Korukonda et al., 2026). We assessed the impact of hTyr-induced pigmentation on LC-associated behaviors, including novelty-induced anxiety, arousal, and cognition, at 1 week post-viral infusion. Consistent with the LC hyperactivity phenotype, we found that bilateral hTyr-injected mice were significantly more reactive in the novelty-suppressed feeding task, a novelty-induced stress paradigm commonly used to model anxiety that is sensitive to changes in LC-NE transmission (Lustberg et al., 2020). hTyr-expressing mice took significantly longer than EYFP controls to consume food in the novelty-suppressed feeding task (t_(24)_ = 2.359, p < 0.05) (**Fig. 5a**). Importantly, hTyr- and EYFP-injected mice displayed no differences in latency to eat in the home cage (data not shown), indicating the experimental group’s increased latency to eat during the task resulted from novelty-induced stress behavior rather than changes in satiety state. There were no differences in arousal, as measured by latency to fall asleep following gentle handling (**Fig. 5b**) or associative memory, as measured by freezing in a footshock-associated context (**Fig. 5c**).

**Fig. 5.**
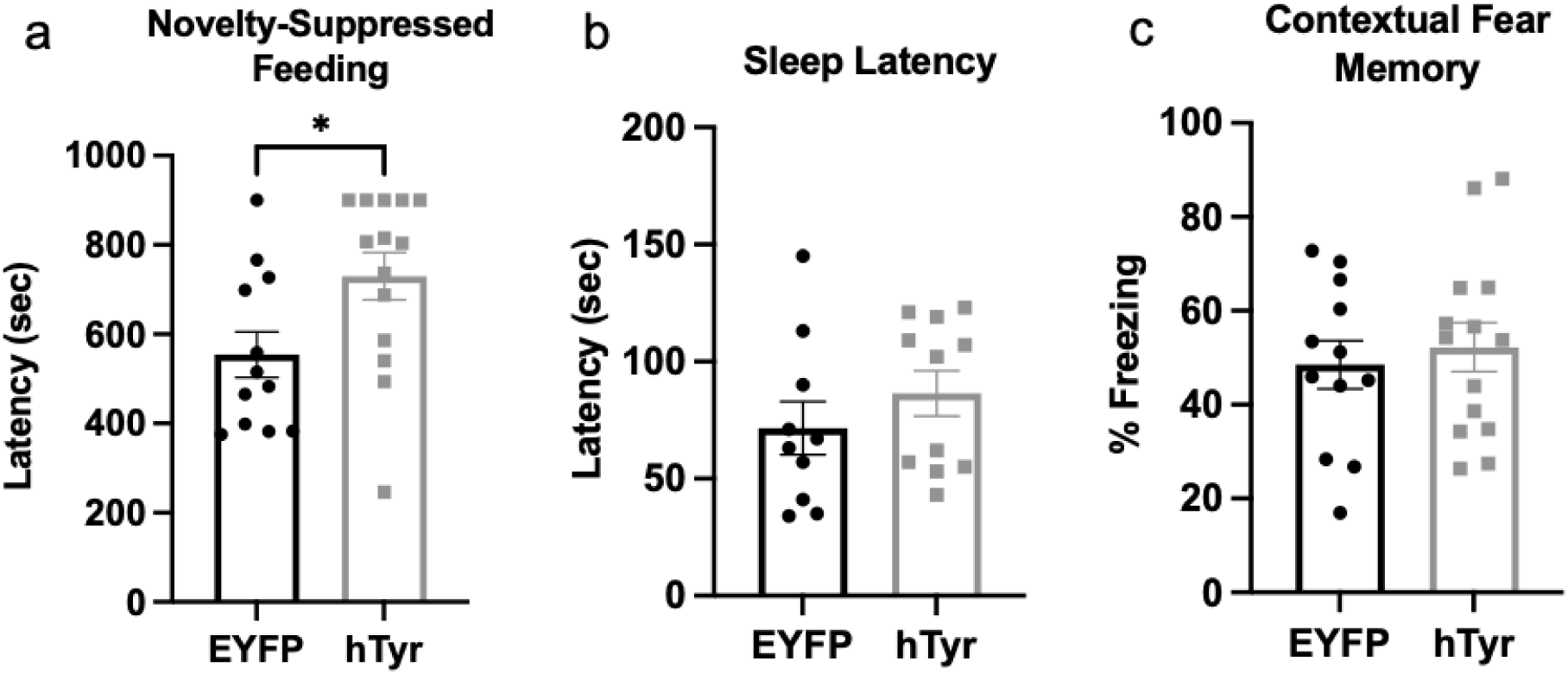
hTyr-induced NM increases anxiety-like behavior but not arousal or contextual fear memory at 1 week. TH-Cre mice received bilateral stereotaxic infusion of AAV-DIO-hTyr or AAV-DIO-EYFP into the LC, and behavior was assessed 1 week later. Shown is mean ± SEM (**a**) latency to bite the food pellet in the novelty-suppressed feeding test, (**b**) latency to fall asleep in the home cage following gentle handling, and (**c**) freezing behavior in the contextual conditioning fear assay. N=12-14 per group. *p<0.05.

### NM alters the LC transcriptome at 1 week

To determine how the LC responded to NM accumulation, we assessed the LC-NE mRNA expression using TRAP. TH-Cre+, *Slc6a2*-*eGFP*/*Rpl10a+* mice received bilateral hTyr or EYFP virus, and brainstem tissue was collected 1 week later. Differential expression (DE) analysis was first performed comparing LC (TRAP) and input (whole-tissue) RNA samples for enrichment of known LC marker genes to verify the method (Mulvey et al., 2018; Iannitelli et al., 2023b). As expected, TRAP fractions were enriched for genes including *Slc6a2*, *Th*, and *Dbh* with log2 fold-change (logFC) values ranging from 2-5. We then sought to examine NM-induced effects on LC-specific and whole hindbrain translatomes by performing DE analyses between genotypes (**Fig. 6a**). The full results of these DE analyses are in **Supplemental Table 3**.

**Fig. 6.**
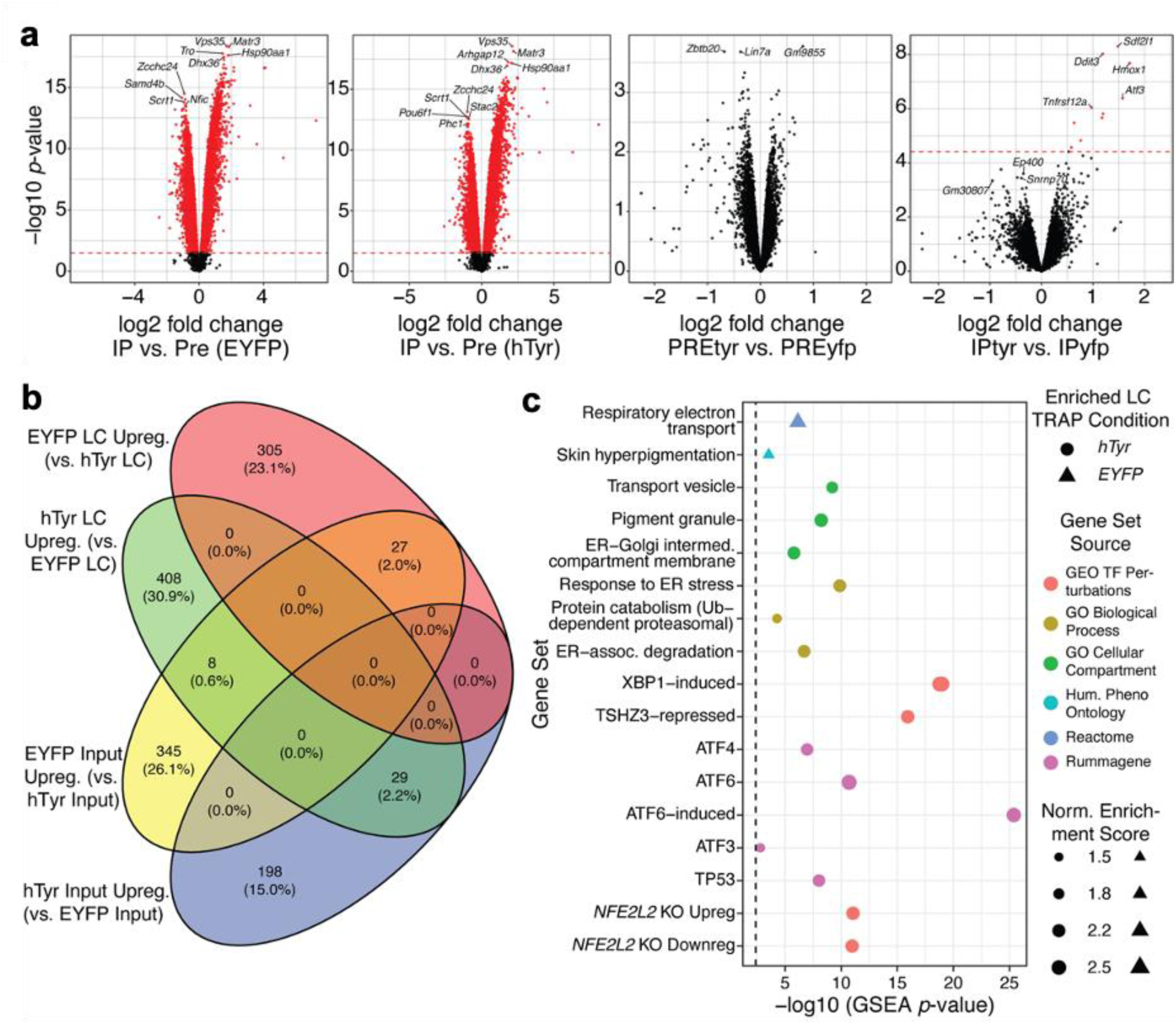
TRAP-seq reveals LC-restricted transcriptional responses to hTyr expression indicative of endoplasmic reticulum stress. TH-Cre+, *Slc6a2*-*eGFP*/*Rpl10a+* mice received bilateral stereotaxic infusion of AAV-DIO-hTyr or AAV-DIO-EYFP into the LC, and gene expression was assessed 1 week later by translating ribosome affinity purification. (**a**) Volcano plots of differential expression (DE) comparisons from TRAP-seq experiments. Red lines demarcate the FDR threshold (0.05) for considering DE significant. On the left, comparisons between IP (TRAP) and Pre-IP (“input”) fractions are shown for mice transduced with EYFP or hTyr, illustrating the magnitude of TRAP enrichment/depletion is similar between the two treatments and that the top-enriched genes (labeled points) are largely consistent across treatments. On the right are shown comparisons of the Pre-IP or IP across treatments, illustrating that hTyr does not substantially change the surrounding tissue transcriptome relative to EYFP, while specifically inducing a small set of proteostasis genes in the LC. (**b**) Venn diagram comparing the number of genes upregulated at a nominal p<0.05 in the EYFP or hTyr condition for each tissue fraction. Note that the vast majority of hTyr-LC upregulated genes are not hTyr-induced in the input sample (29/408 genes), indicating the transcriptional effects of hTyr are LC-specific. (**c**) Select GSEA results from DE analysis between hTyr IP and EYFP IP. The upper half of the plot illustrates ontology and pathway terms, while the lower half illustrates TFs whose target genes are enriched in DE patterns. The size of each point indicates the degree of DE enrichment for the term, the color indicates the source for the term, and the shape of the point indicates whether the term was enriched in EYFP- or hTyr-upregulated genes. As pathways and TF-target sets were tested separately, the more stringent of the two FDR<0.05 thresholds is shown as a dashed vertical line.

Notably, we observed 10 significant (FDR < 0.05) DE genes (DEGs) between hTyr LC and EYFP LC, but none between input fractions from the two conditions, indicating the specificity of our methods for capturing LC mRNA. At nominal significance (p < 0.05), we identified 623 DEGs between hTyr input and EYFP input and 813 DEGs between hTyr LC and EYFP LC (**Fig. 6a**); these nominal DEGs were also mostly exclusive to the respective RNA fractions, further supporting LC-specific effects. The top individual genes upregulated in hTyr LC were *Hmox1*, *Sdf2l1*, *Ddit3* (aka GADD34), and *Atf3*; while no genes were significantly downregulated in hTyr LC, those nearest to significance included *Ep400* (FDR=0.144). To understand the functional potential of hTyr-induced gene expression changes in LC, we then utilized GSEA (see *Methods*) with pathways from MsigDB to identify ontologies, cell types, and pathways enriched in genes DE in LC. Top terms enriched in hTyr-LC upregulated genes included pigmentation and multiple facets of ER stress response. (**Fig. 6b** and **Supplemental Table 4**).

To identify transcriptional regulators downstream of hTyr-induced NM aggregation, we also performed GSEA (see *Methods*) on sets of transcription factor (TF) target genes cataloged in several other resources. The top 20 sets were comprised almost exclusively of various annotated target genes of TFs involved in unfolded protein response (UPR), namely *Atf6* and *Xbp1*. Neither of these TFs were differentially expressed between EYFP-LC and hTyr-LC, however. To decipher possible means by which the targets of these TFs could be robustly upregulated without change in TF expression, we identified 61 proteins which have "high confidence" interactions with *Atf6 and/*or *Xbp1* in mouse in StringDB v12.0. Among these interacting proteins, three were significantly DE (all upregulated) in hTyr-LC relative to EYFP-LC: *Ddit3*, *Atf3*, and *Ppp1r15a*. Interestingly, all three of these genes are also implicated in UPR or other forms of endoplasmic reticulum stress, and two are themselves TFs (Bernstein et al., 2011; Chalour et al., 2018; El Manaa et al., 2021).

Given our findings of UPR in the hTyr LC, along with a substantial body of research implicating this pathway in the detrimental effects of neuropathological aggregates, we next sought to characterize whether AD pathology induced corresponding changes in TF expression in non-NM neurons. Interestingly, the five aforementioned TFs were not DE between bulk RNA from AD and control postmortem brain regions (see *Acknowledgments*), nor were they consistently differentially expressed at the level of cortical cell types identified in a large single-nucleus study of AD (Mathys et al., 2023). To further examine this relationship, we performed the same TF-target GSEA analysis as above, using entorhinal excitatory neuron DE results between early AD pathology/controls and between late/early AD pathology (Mathys et al., 2019). Strikingly, target sets with significant enrichment in early vs. no AD pathology DEGs were negatively enriched (i.e., more highly expressed in controls than early AD), while they were upregulated in late AD relative to early AD (**Supplemental Table 3**). These findings suggest that the UPR pathway induced by NM in the LC is distinct from that induced in neurons more broadly by early pathological aggregates.

### Persistent NM accumulation results in neurodegeneration and neuroinflammation in the LC and projection fields

To assess the consequences of prolonged pigment burden in the LC, TH-Cre+ and TH-Cre- mice received AAV-DIO-hTyr and were assessed 6 weeks following bilateral viral infusion. At this time point, we observed a catastrophic loss of LC cell bodies, as measured by TH immunoreactivity (t_(3)_ = 30.39, p < 0.0001) (**Fig. 7b-f**) and Nissl (t_(3)_ = 25.58, p < 0.0001) (**Fig. 7b-g**). This neurodegeneration was accompanied by profound increases in Iba-1 (t_(6)_ = 5.04, p = 0.001) (**Fig. 7b, d, g**) and GFAP (t_(5)_ = 3.59, p = 0.016) (**Fig. 7c, e, h**) immunoreactivity, indicating a local neuroinflammatory response. Strikingly, reactive astrocytes had completely infiltrated and filled the space vacated by dying NE neurons (**Fig. 7c**). Although only a few LC neurons remained at this time point, many darkly pigmented granules were detectable in the LC region and were also visible beyond its typical confines (**Fig. 7a**), indicating that NM persists and may spread to adjacent tissue following the death of their host neurons of origin. The degeneration of LC cell bodies was associated with a near-complete loss of NET+ fibers in the hippocampus (**Fig. 7i, l, u**) and prelimbic cortex (**Fig. 7**). There was also an increase of Iba-1 immunoreactivity in the hippocampus and an increase of both Iba-1 and GFAP in the prelimbic cortex.

**Fig. 7.**
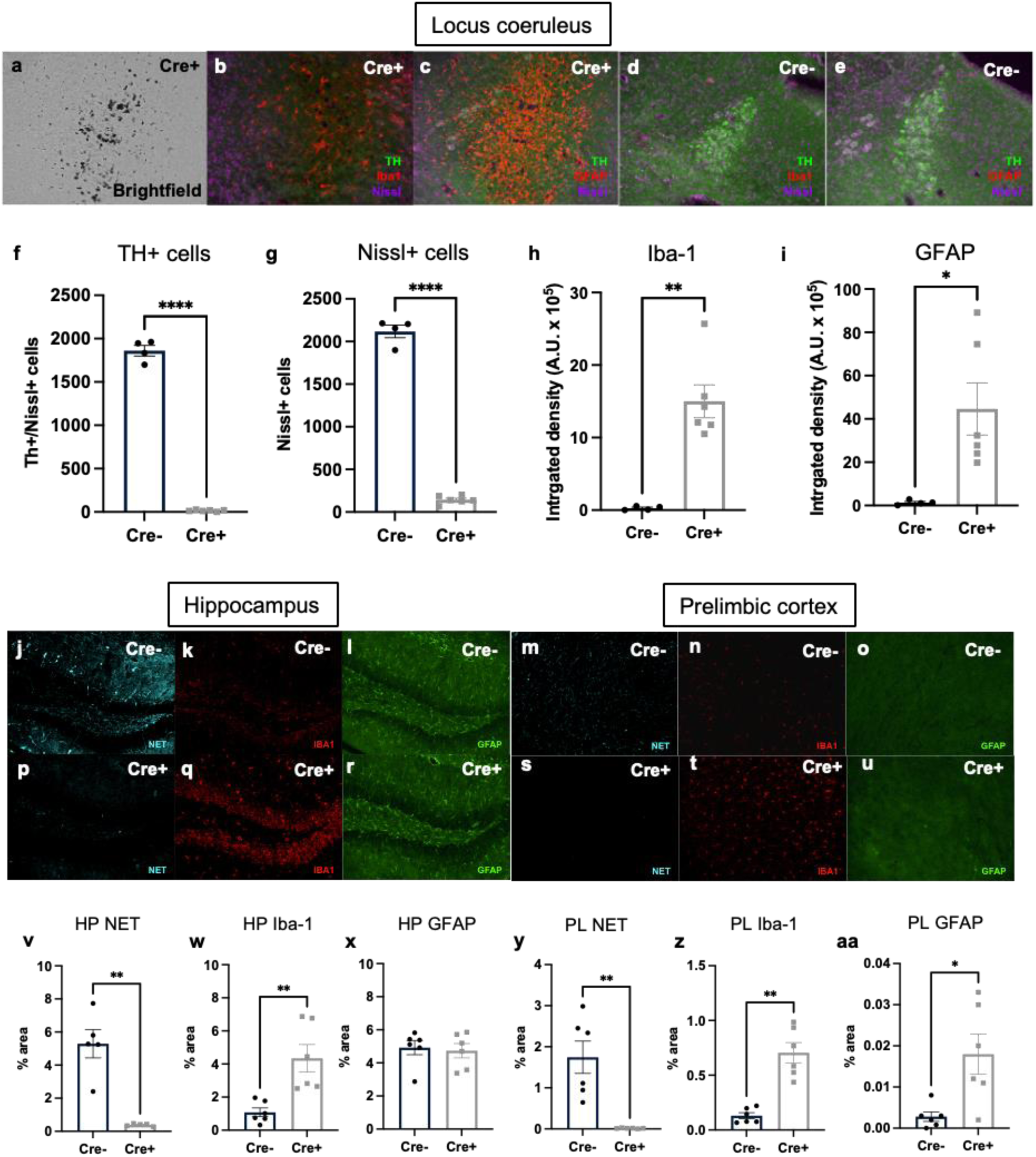
Prolonged NM accumulation results in LC degeneration and neuroinflammation. TH-Cre+ and TH-Cre- mice received bilateral stereotaxic infusion of AAV-DIO-hTyr into the LC and aged for 6 weeks. Shown are representative images of (**a**) brightfield NM, (**b**, **d**) TH + Iba-1 + Nissl, (**c**, **e**) TH + GFAP + Nissl. Quantification (mean ± SEM) is shown for (**f**) TH+/Nissl+ cells, (**g**) Nissl+ cells, (**h**) Iba-1, and (**i**) GFAP. Representative immunofluorescent images are shown for (**j**, **m**) NET, (**k**, **n**) Iba-1 and (**l**, **o**) in the hippocampus and (**p**-**u**) prelimbic cortex. Quantification (mean ± SEM) is shown in **u**-**aa**. GFAP, hippocampus; PL, prelimbic cortex. Immunofluorescence images were acquired at 20X. N = 4-6 per group. *p<0.05, **p<0.01, ****p<0.0001.

To more fully characterize the ultrastructural composition and distribution of NM in this model over time, we performed electron microscopy on LC tissue. As previously reported for endogenous human NM, the hTyr-induced NM aggregates were characterized as clusters made up of irregularly shaped pigment granules of various electron density and sizes, as well as lipid droplets. Overall, the clusters of NM were found in neuronal cell bodies (**Fig. 8a, c**), large membrane-delimited neuropil aggregates (**Fig. 8b, d, f**), and glial cells (**Fig. 8e, g**). To determine the phenotype of the neuronal profiles that contained NM, some sections were immunostained for TH to mark LC-NE neurons, while others were labeled for either the astrocytic marker GFAP or the microglia marker Iba-1. As shown in **Fig. 8c**, some TH-positive cell bodies and dendritic profiles were enriched in NM, confirming that NM is found within noradrenergic neurons in the LC. Regarding glial expression, our observations revealed clear examples of NM aggregates within Iba-1-positive microglia cell bodies (**Fig. 8e, g**), but not in GFAP-positive astrocytes (**Fig. 8f**). However, in some instances, GFAP-positive processes were closely apposed to NM-containing profiles (**Fig. 8f**). We speculated that the persistence of NM in the LC region after many of the NE neurons were lost could be explained by glial engulfment. To test this idea, we counted the total number of neuronal and glial profiles that contained NM in the tissue and calculated the ratio of glia:neuron profile labeling. As shown in **Fig. 8h**, this ratio increased with the length of the post-viral infusion period, from ∼5% at 1 week to ∼25% at 10 weeks. These data suggest that NM initially accumulates in LC-NE neurons but gradually invades the LC neuropil following release upon neuronal death, where it is engulfed by microglia. NM was absent in controls (i.e. mice that received EYFP virus or no virus; **Fig. 8h** and data not shown). With the use of serial EM images (**Figs. 8i-k**, and **Figs. 8m-p**) and the 3D reconstruction approach (**Figs. 8l, q**), we obtained 3D models of microglia cells (**Fig. 8q**) and their process (**Fig. 8l**) that allowed the spatial visualization of the NM clusters **(Fig. 8l)** and the complete 3D visualization of microglia cells. These 3D images showed that a large volume of the microglia cytoplasm is occupied by multiple NM clusters and other cytoplasm inclusions of degenerated axonal profiles, and that some processes of GFAP-positive astrocytes are partially covering the cell surface (**Figs. 8l, q; Supplemental Video 1**). One limitation of our analysis is that because much of the NM was associated with degenerating neuronal elements (e.g. dendrites) at later time points, it was difficult to discern “intracellular” vs “extracellular” localization with confidence. Given that most of the LC-NE neurons were dead by 6 weeks, we assume that much of it was no longer associated with intact cells and was accessible to invading glial at these time points.

**Fig. 8.**
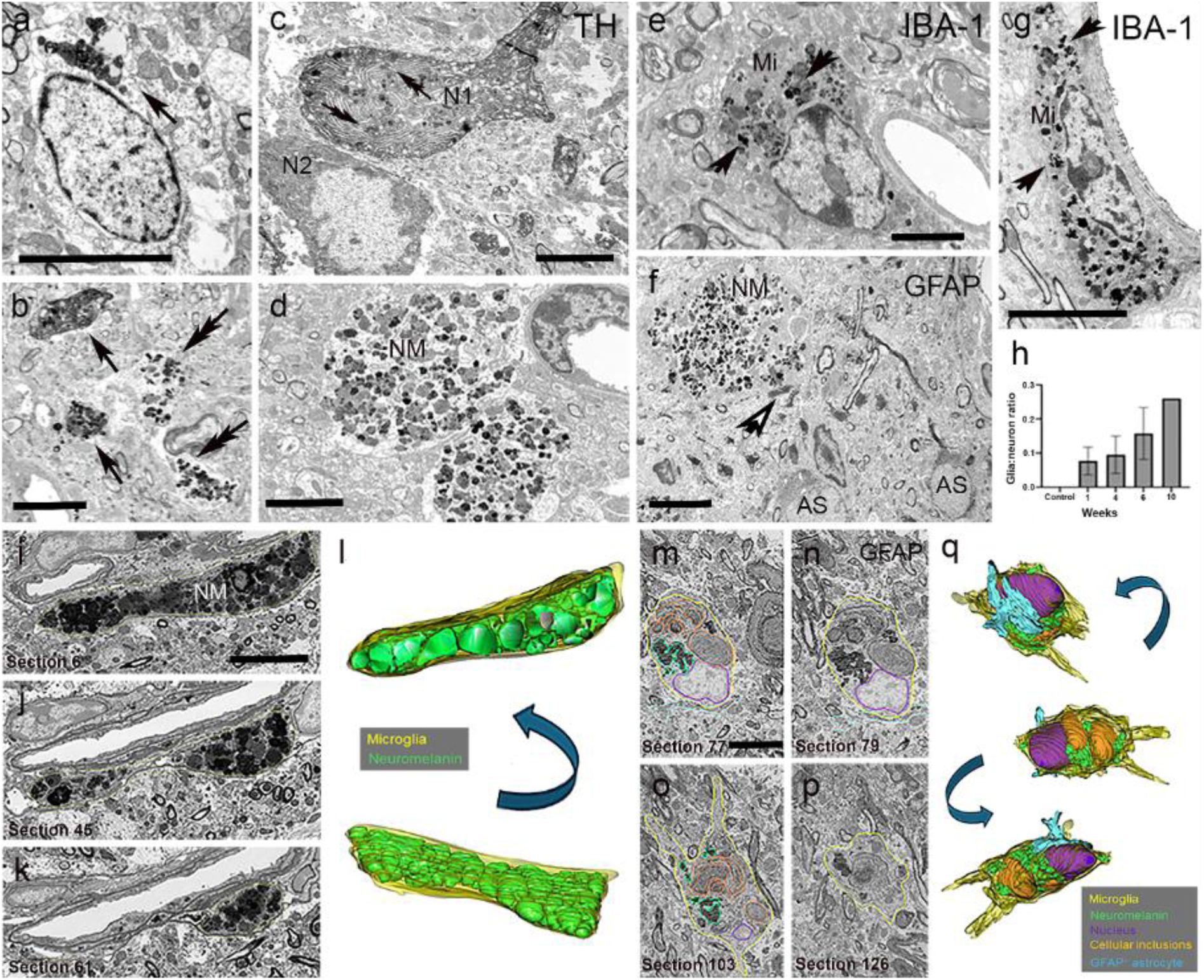
Transmission 2D EM analysis and 3D ultrastructural reconstruction of neuromelanin (NM) granules in the mouse LC. TH-Cre mice received bilateral stereotaxic infusion of AAV-DIO-hTyr or EYFP control into the LC, and electron microscopy was performed 1-10 weeks later. (**a**) Cluster of NM in the cell body of a LC neuron (arrow, 1 week hTyr), (**b, c**) LC tissue (1 week hTyr) immunostained for TH showing NM expression in TH-positive dendritic profiles (arrows in **b**) and a neuronal cell body (N1 in **c**). In **b**, the double arrows indicate NM-containing TH-negative dendrites, while in **c**, they point at intracellular NM granules. Note in **c**, a TH-positive LC neuronal perikaryon (N2) that does not contain NM. (**d**) Large neuropil structures heavily packed with NM granules in the LC (6 weeks hTyr). (**e, g**) Examples of immunoperoxidase-stained IBA-1-containing microglia (Mi) cell bodies enriched in NM aggregates (arrows) in the LC of a 6 week hTyr mouse. (**f**) GFAP-positive astrocytic profiles (AS) in the close vicinity of a NM-containing neuropil element (6 weeks hTyr). Note the lack of NM in AS cell bodies, and the close apposition between an AS process (open arrow in **f**) and the NM-containing element. (**h**) The ratio of NM-containing glia:neuron profiles in the LC of control (no hTyr: n=3) and experimental (hTyr 1 week: n=3; 4 weeks: n=2; 6 weeks: n=4; 10 weeks: n=1) mice. (**i-k, m-p**) Serial ultrastructural images of NM clusters (**i-k**) and a putative microglia cell body profile **(m-p)** from the LC of a 6 week hTyr mouse used in *Reconstruct* (NIH) to create their respective 3D models (**l, q**). Different rotated images of the 3D reconstructed models showing the NM clusters (**l**) and the intracellular distribution of NM, cytoplasm inclusions of degenerated axonal profiles and nucleus in the microglia (**q**). The tissue used for the 3D analysis of the microglia was immunostained with GFAP and allowed for the analysis of the spatial interaction between the microglia and GFAP-positive astrocytic processes. A total of 83 and 139 serial images, respectively, were used in the 3D reconstruction of the NM clusters (**l**) and the microglia (**q**). Scale bars in **a-g, i** (applies to **j, k**) and **m** (applies to **n-p**) = 5µm. Abbreviations: Mi=Microglia; NM=Neuromelanin; N=Neuron; AS=Astrocyte. Scale bar value: A: 5 mm; **b**, **c**, **f**: 2 mm; **d**, **e**, **g**: 3 mm.

### NM-induced LC neuron death is cell-autonomous

There is evidence for both intracellular and extracellular NM toxicity (Iannitelli and Weinshenker, 2023). For example, hTyr-induced NM accumulation in the rat SNc induces a general failure of proteostasis, while enhancement of lysosomal-mediated proteolysis reduces intracellular neuromelanin and prevents DA neuron degeneration (Carballo-Carbajal et al., 2019). On the other hand, there is evidence that NM released from dying neurons can contribute to the degeneration of neighboring cells; application of NM to neuronal cultures or infusion of purified NM into the rat SNc kills resident DA neurons (Zhang et al., 2011). Indeed, we observed NM granules in the LC region even after all the TH+ neurons were gone (**Fig. 7**). Because infusion of hTyr virus into the LC of TH-Cre mice results in NM production and death in nearly all LC neurons, we could not distinguish intracellular (i.e. cell-autonomous) from extracellular (i.e. cell non-autonomous) toxicity.

To address this question, we infused EYFP virus into one hemisphere of the LC and hTyr virus into the other hemisphere of the LC of Pdyn-Cre, which express Cre in only a subset of LC neurons, and examined EYFP, TH, and NM expression 10 weeks later (representative images shown in **Fig. 9a, b**). We identified 10 weeks as a point long after NM kills LC neurons to provide NM released from dying cells time to damage adjacent cells. We reasoned that if NM-induced cell death were cell autonomous, then only the LC neurons expressing hTyr would die, whereas if NM was being released from dying cells and killing neighboring cells, then even LC neurons not expressing Cre would be killed. We compared the “predicted” cell death (i.e. the number of Cre-positive cells on the EYFP side, calculated as the number of TH+ neurons expressing EYFP) to the “actual” cell death (i.e. the difference between the number of TH+ cells on the EYFP side and the number of TH+ cells on the hTyr side), with the null hypothesis being that NM toxicity was strictly cell-autonomous. We found that ∼14% of the TH+ cells on the EYFP side co-expressed EYFP (114 ± 27 out of 828 ± 23) (**Fig. 9c**), and NM was observed on the hTyr side (**Fig. 9b**). EYFP was also expressed in some TH-cells, and NM was observed outside the confines of the LC core, (**Fig. 9a, b**), indicating that endogenous catecholamines are not required for hTyr-induced NM formation. As expected, there were significantly fewer TH+ cells on the hTyr side, (t = 5.30, p = 0.003), indicating neuronal death (**Fig. 9d**. Importantly, paired t-test revealed no significant difference between the predicted and actual degree of cell death on the hTyr side (t = 1.59, p = 0.17) (**Fig. 9e**), suggesting that NM toxicity was confined to the cells that expressed it and did not cause degeneration of neighboring cells after release. Given our electron microscopy results, it is possible that microglial engulfment of extracellular NM protected surviving LC neurons from its toxicity.

**Fig. 9.**
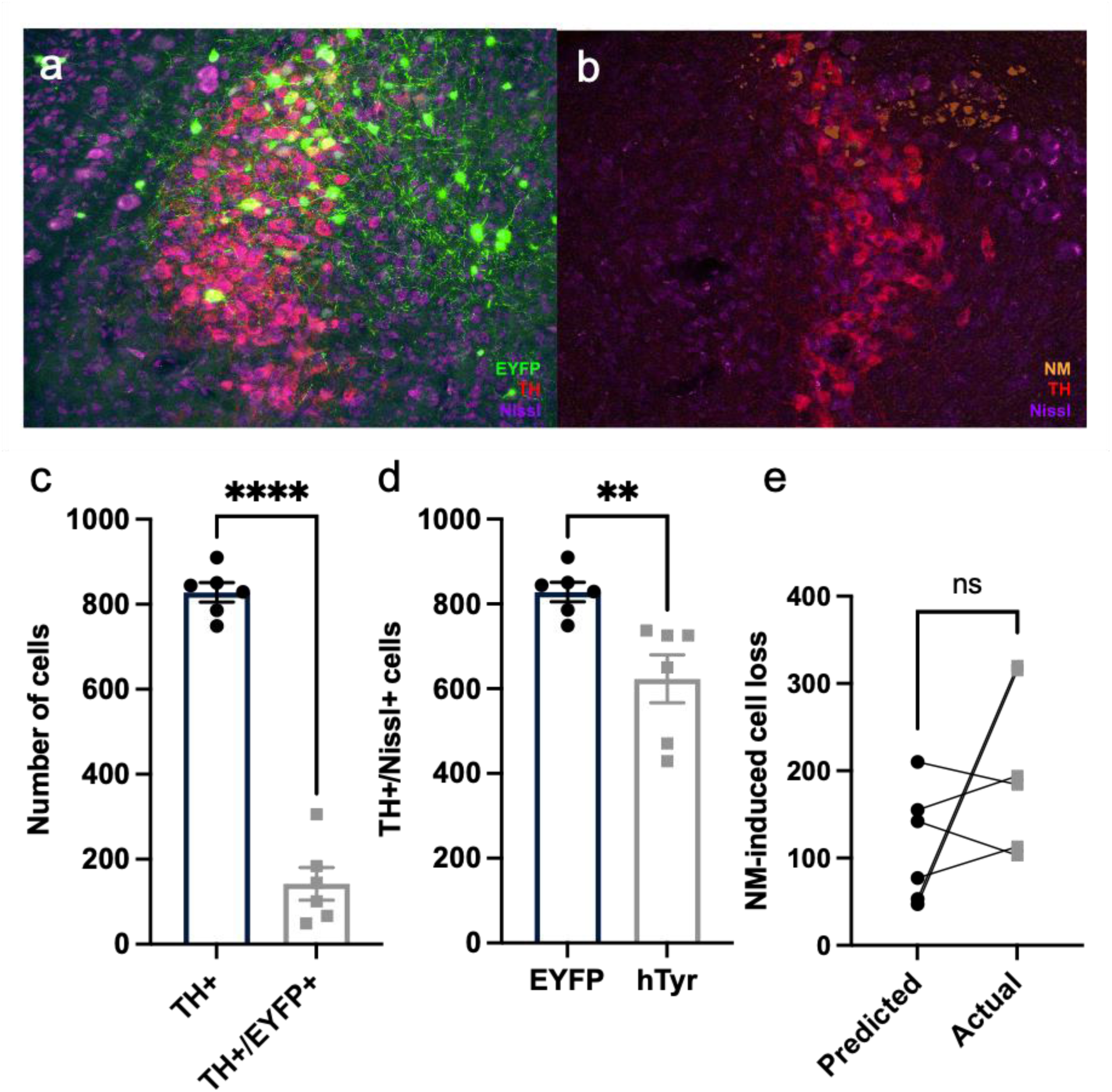
NM-induced cell death in the LC is cell-autonomous. Pdyn-Cre mice received AAV-DIO-hTyr on one side of the LC and AAV-DIO-EYFP on the other side of the LC and assessed 10 weeks later. Shown are representative immunofluorescent images of (**a**) EYFP or (**b**) NM (pseudocolored from brightfield) in combination with TH and Nissl, as well as quantification for (**c**) TH+ vs TH+/EYFP+ cells for the EYFP side, (**d**) TH+/Nissl+ cells, and (**e**) predicted (number of TH+/EYFP+ cells on the EYFP side) vs actual (difference between the number of TH+ cells on the EYFP side and the number of TH+ cells on the hTyr side) TH+/Nissl+ cell loss. Immunofluorescence images were acquired at 20X. N = 6 per group. **p<0.01, ****p<0.0001.

### NM-mediated neurodegeneration has no behavioral effects at 6 weeks

Despite the catastrophic loss of LC neurons at 6 weeks in mice with bilateral NM accumulation, we did not observe any significant differences in behavior compared to mice that received EYFP virus. Notably, novelty-induced anxiety, which was increased in hTyr mice at 1-week, did not differ statistically between groups at 6 weeks (**Fig. 10a**). Pigment-expressing mice also displayed no differences in sleep latency (**Fig. 10b**) or contextual fear conditioning (**Fig. 10c**).

**Fig. 10.**
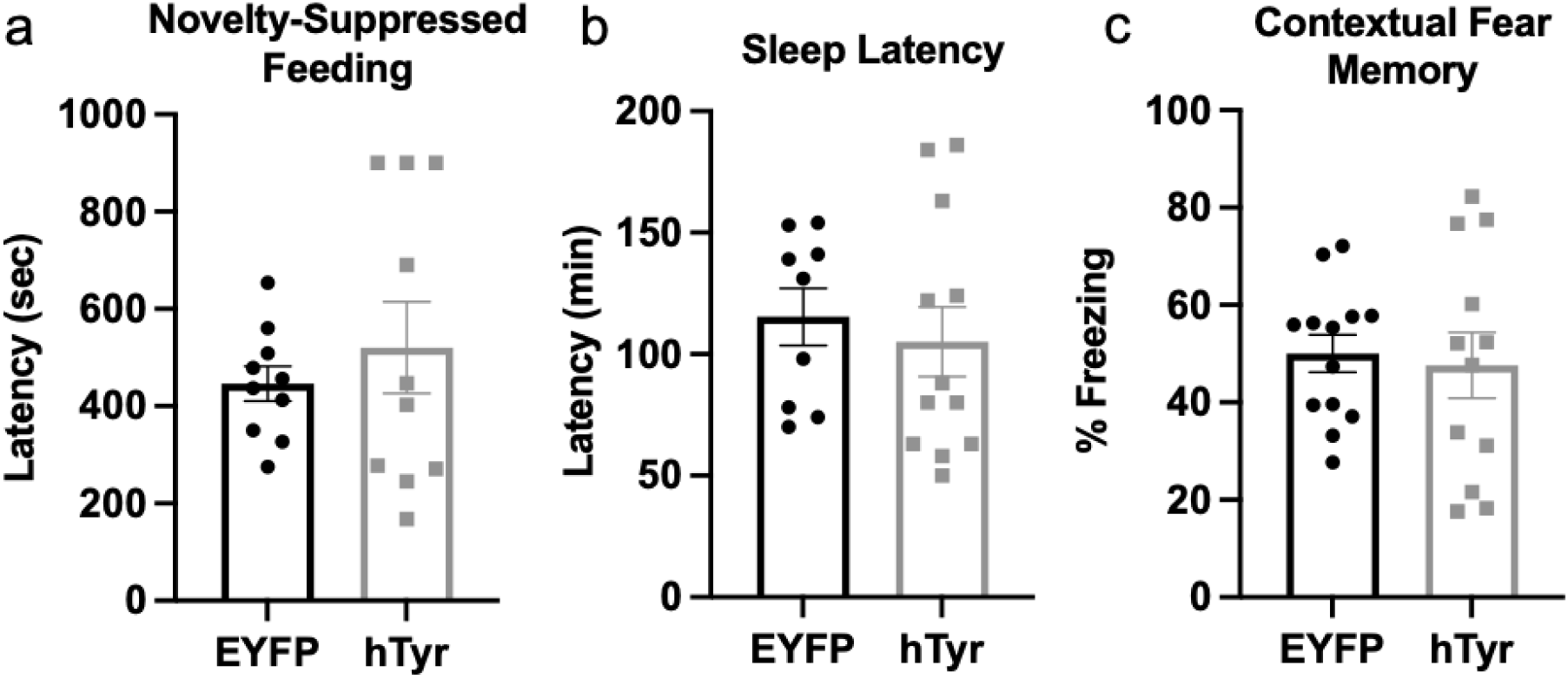
NM accumulation and subsequent neurodegeneration have no effect on behavior at 6 weeks. TH-Cre mice received bilateral stereotaxic infusion of AAV-DIO-hTyr or AAV-DIO-EYFP into the LC, and behavior was assessed 6 weeks later. Shown is mean SEM (**a**) latency to bite the food in the novelty-suppressed feeding test, (**b**) latency to fall asleep following gentle handling, and (**c**) % time freezing in contextual fear conditioning. N=10-13 per group.

## Discussion

The present study characterized the impact of hTyr-induced NM in the LC of mice, allowing the assessment of causal relationships between NM accumulation in noradrenergic neurons, LC-NE integrity and activity, and the development of non-motor symptoms reminiscent of prodromal AD and PD in humans. The expression of NM driven by viral delivery of hTyr is consistent with prior work focused on the SN (Carballo-Carbajal et al., 2019). In our model, neuronal dysfunction was evident as early as 1 week post-infusion. Even at this early time point, we found reduced TH immunoreactivity in the pons, with a trend towards loss of NET immunoreactivity in the hippocampus of hTyr mice compared to controls, suggesting loss of fiber/terminal integrity, but no difference in the number of noradrenergic or total LC cell bodies. These findings are consistent with other reports of early LC axon damage in neurodegenerative diseases (Weinshenker, 2018; Doppler et al., 2021b; Gilvesy et al., 2022), as well as a loss of catecholamine identity following hTyr/NM expression in the SN of rats (Carballo-Carbajal et al., 2019). We also observed increased Iba-1 signal in terminal fields, and microglia have been shown to phagocytose LC axons in an APP mouse model of AD (Meyer et al., 2025). Concurrent with fiber loss was depletion of NE and its primary metabolite MHPG in the hippocampus and PFC. The MHPG:NE ratio, a measure of NE turnover, was elevated in projection regions, suggesting increased release from surviving LC fibers described in rodents (Hallman and Jonsson, 1984; Iannitelli et al., 2022) and humans (Francis et al., 1985; Raskind et al., 1999; Jacobs et al., 2021) with LC neurodegeneration. Using slice electrophysiology, we found that LC-NE neurons containing NM were hyperactive, including increases in both spontaneous and evoked firing. Excessive NE transmission is a common feature of compromised LC neurons, and can occur at the cell body, terminal, and/or postsynaptic receptor level, depending on the nature of the insult (Butkovich et al., 2020; Kelly et al., 2021; Iannitelli et al., 2023b; Kelberman et al., 2023; Wang et al., 2025; Korukonda et al., 2026). We speculate that this hyperactivity is not a direct result of pathology per se, but rather a compensatory response to damage in an effort to maintain normal NE transmission, reminiscent of compensatory LC hyperactivity following chronic inhibition by opioids (Maldonado, 1997; Mazei-Robison and Nestler, 2012).

A NE hyperactivity phenotype is congruent with the LC dysfunction seen in prodromal AD and PD (Weinshenker, 2018). NM-expressing mice displayed elevated novelty-induced anxiety-like behavior at 1 week, indicating that the compensatory LC-NE hyperactivity has behavioral consequences (Lustberg et al., 2020; Kelberman et al., 2022; Iannitelli et al., 2023b; Korukonda et al., 2026). The disappearance of the anxiety phenotype at later time points when the LC was completely degenerated is consistent with this interpretation. It was somewhat surprising that there was no effect on this hyperactivity on sleep latency, as NE transmission promotes arousal (Carter et al., 2010; Porter-Stransky et al., 2019; Poe et al., 2020). Sleep is a complex process controlled by many brain regions and neurotransmitters, and it is possible that other circuits are compensating in response to the LC hyperactivity in this case (Korukonda et al., 2026).

In the rat hTyr model, NM accumulation in the SNc induces a general proteostasis failure in these neurons, and enhancing proteostasis via overexpression of transcription factor EF (TFEB) attenuates NM-induced degeneration (Carballo-Carbajal et al., 2019). Here, RNA-seq of the hTyr-expressing LC revealed large impacts of induced NM production on the transcriptome of LC neurons, markedly converging on UPR pathways. Notably, target genes of UPR TFs *Atf6* and *Xbp1* were the most robustly upregulated in hTyr-LC, though these two TFs themselves did not show a change in expression compared to EYFP-LC. Rather, several TFs which interact with *Atf6* and/or *Xbp1* and which are also members of the UPR pathway were upregulated, suggesting that NM induces *Atf6*- and *Xbp1*-associated genes through activity of interacting TFs (*Ddit3*, *Atf3*). Moreover, we found that the targets of these same TFs are *downregulated* in excitatory neurons of the entorhinal cortex in early AD (Mathys et al., 2019), suggesting the UPR pathway induced by NM may counter neuronal UPR pathway effects of early proteinopathy. Notably, *XBP1* overexpression enables human neuron survival in the presence of ordinarily cell-fatal amounts of Aβ (Duran-Aniotz et al., 2023), suggesting that induction of these TFs in LC in our model represents an adaptive neuronal response to the presence of NM. *Atf6* and *Xbp1*, whose targets appeared to be upregulated by NM, are attributed to the “adaptive”, rather than pro-apoptotic, arm of the UPR (Hetz et al., 2020). Consistent with the observation that LC cell bodies can persist for years with high pTau burden (Braak et al., 2011), our findings suggest a potential mechanism by which NM acts as *both* a cellular insult *and* a neuroprotectant. For example, in responding to accumulating NM, the LC upregulates *Atf3/Atf6/Xbp1/Ddit3* or their targets, allowing the cell to tolerate the effects of NM (to a limited extent) while simultaneously inducing a molecular state that is protective against the toxic effects of protein aggregates found in neurodegenerative diseases.

Substantial cell death and neuroinflammation in the LC were evident by 6 weeks post-viral infusion. Transgenic expression of hTyr driven by the TH promoter similarly provokes LC cell death in mice, with animals displaying a partial loss of LC neurons by 1 month of age and near-total LC death by 3-4 months (Laguna et al., 2024). By contrast, DA neuron death in the SN was delayed and milder in this model, similar to what is observed in clinical PD (Braak et al., 2003; Zarow et al., 2003; Laguna et al., 2024). Dense NM persisted in the region following LC degeneration, suggesting that these granules had been released by dying neurons and either remained in the extracellular space or were taken up by other cells. At the gross immunofluorescent level, there was abundant astrocytic and microglial inflammation, but while electron microscopic analysis showed many astrocyte processes closely apposed to NM granules, only microglia appeared to engulf the NM. These findings are aligned with evidence supporting extracellular NM-induced activation of microglia that has been reported across human postmortem tissue, animal models, and cell culture (Zecca et al., 2003; Zecca et al., 2008a; Zhang et al., 2011). Neuroinflammation is a well-appreciated aspect of AD and PD that is influenced by the LC-NE system (Chalermpalanupap et al., 2013; Feinstein et al., 2016; Giorgi et al., 2020; Mercan and Heneka, 2022; Tansey et al., 2022). Our data suggest that the release of pigmentation from degenerating LC neurons contributes to the neuroimmune response. Despite evidence that extracellular NM can be toxic to catecholamine neurons (Zhang et al., 2011), the experiment using the Pdyn-Cre mice that expressed hTyr in only a subset of LC neurons suggests that cell death is primarily cell-autonomous in our model. One possibility is that microglial engulfment of NM released from dying neurons protected neighboring cells, but this remains to be tested. There were no differences between groups in anxiety-like behavior, arousal, or contextual fear memory at the 6-week time point. The anxiogenic novelty-suppressed feeding phenotype observed at 1 week was absent at 6 weeks, which may be explained by a near-total loss of LC-NE, which is required for this behavior (Lustberg et al., 2020).

The present study provides novel insights into the consequences of NM accumulation in rodent LC-NE neurons but also has some limitations. An important aspect of our approach is that viral hTyr overexpression drives extremely rapid NM accumulation over the time course of weeks, as compared to natural NM, which accumulates in humans over decades. This condensed timescale may dramatically accelerate the ability of NM to overwhelm cellular machinery and lead to cell death. In this respect, our approach shares the same limitation as other rodent models of neurodegenerative disease. Nevertheless, the rate of NM accumulation can increase under some conditions. For example, people with post-traumatic stress disorder, which is associated with LC hyperactivity (Naegeli et al., 2018), have a higher LC NM-MRI signal (McCall et al., 2024), and there is ample evidence that the same processes contribute to LC neuron dysregulation and demise in AD and PD. Indeed, LC neurons with the highest NM content are disproportionally lost in PD (Mann and Yates, 1983b). It also appears that LC neurons are more vulnerable to NM-induced toxicity than DA neurons (Laguna et al., 2024). Ultimately, this model of pigment induction may be critical for uncovering the role NM plays in the early vulnerability of LC neurons in AD and PD.

## Author Contributions

L.H. designed and performed researched, analyzed data, wrote the first draft of the manuscript, and edited the manuscript. B.M. designed research and analyzed data. H.E.B. performed research and analyzed data. M.T. designed and performed research. A.K. designed and performed research. L.C.L. designed and performed research. A.L.S. performed research and analyzed data. J-F.P. performed research and analyzed data. R.V. performed research and analyzed data. X.C. performed research and analyzed data. K.H. performed research and analyzed data. J.J.Y. designed research. D.E.H. designed research, analyzed data, and wrote the first draft of the manuscript. S.K. designed research. A.S. performed research and analyzed data. S.A.S. designed research and analyzed data. K.M. analyzed data. K.Y. designed research. J.D.D. designed research and contributed analytic tools. K.E.M. performed research and analyzed data. Y.S. designed research, analyzed data, wrote first draft of the manuscript, and edited the manuscript. M.J.B. designed and performed research, analyzed data, wrote the first draft of the manuscript, and edited the manuscript. D.W. designed research, analyzed data, and wrote the manuscript. A.F.I. designed and performed research, analyzed data, wrote the first draft of the manuscript, and edited the manuscript.

## Acknowledgements

This work was supported by the National Institute on Aging (AG079199 to D.W. and M.J.B., AG061175 to D.W., K.Y., and S.K., AG085933 to K.M., AG079620 to H.E.B., AG081046 to A.K.), the National Institute of Neurological Disorders and Stroke (NS129168 to A.F.I. and NS135830 to M.J.B. and D.W.), the National Institute of Environmental Health Sciences (ES12870 to A.F.I.), the National Institute of Mental Health (MH015330 to B.M.), and the NIH/ORIP Emory Primate Center (OD011132 to Y.S.). This study was supported in part by the Emory HPLC Bioanalytical Core, which is subsidized by the Emory University School of Medicine and is one of the Emory Integrated Core Facilities, and the Emory Viral Vector Core, which is subsidized by the Emory Center for Neurodegenerative Disease. The material used in the 3D electron microscopy analysis was prepared at the Multiscale Microscopy Core, a member of the OHSU University Shared Resource Cores RRID:SCR_009969. Additional support was provided by the Georgia Clinical & Translational Science Alliance of the National Institutes of Health under Award Number UL1TR002378. The results published here are in whole or in part based on data obtained from Agora, a platform initially developed by the NIA-funded AMP-AD consortium that shares evidence in support of AD target discovery. Agora is available at: doi:10.57718/agora.adknowledgeportal.

## References

Bankhead P, Loughrey MB, Fernandez JA, Dombrowski Y, McArt DG, Dunne PD, McQuaid S, Gray RT, Murray LJ, Coleman HG, James JA, Salto-Tellez M, Hamilton PW (2017) QuPath: Open source software for digital pathology image analysis. Sci Rep 7:16878.

Barden H, Levine S (1983) Histochemical observations on rodent brain melanin. Brain Res Bull 10:847–851.

Bernstein AI, Garrison SP, Zambetti GP, O’Malley KL (2011) 6-OHDA generated ROS induces DNA damage and p53- and PUMA-dependent cell death. Mol Neurodegener 6:2.

Bondareff W, Mountjoy CQ, Roth M (1982) Loss of neurons of origin of the adrenergic projection to cerebral cortex (nucleus locus ceruleus) in senile dementia. Neurology 32:164–168.

Braak E, Sandmann-Keil D, Rub U, Gai WP, de Vos RA, Steur EN, Arai K, Braak H (2001) alpha-synuclein immunopositive Parkinson’s disease-related inclusion bodies in lower brain stem nuclei. Acta Neuropathol 101:195–201.

Braak H, Del Tredici K (2011a) Alzheimer’s pathogenesis: is there neuron-to-neuron propagation? Acta Neuropathol 121:589–595.

Braak H, Del Tredici K (2011b) The pathological process underlying Alzheimer’s disease in individuals under thirty. Acta Neuropathol 121:171–181.

Braak H, Thal DR, Ghebremedhin E, Del Tredici K (2011) Stages of the pathologic process in Alzheimer disease: age categories from 1 to 100 years. J Neuropathol Exp Neurol 70:960–969.

Braak H, Del Tredici K, Rub U, de Vos RA, Jansen Steur EN, Braak E (2003) Staging of brain pathology related to sporadic Parkinson’s disease. Neurobiol Aging 24:197–211.

Branch SY, Beckstead MJ (2012) Methamphetamine produces bidirectional, concentration-dependent effects on dopamine neuron excitability and dopamine-mediated synaptic currents. J Neurophysiol 108:802–809.

Bueicheku E, Diez I, Kim CM, Becker JA, Koops EA, Kwong K, Papp KV, Salat DH, Bennett DA, Rentz DM, Sperling RA, Johnson KA, Sepulcre J, Jacobs HIL (2024) Spatiotemporal patterns of locus coeruleus integrity predict cortical tau and cognition. Nat Aging 4:625–637.

Bush WD, Garguilo J, Zucca FA, Albertini A, Zecca L, Edwards GS, Nemanich RJ, Simon JD (2006) The surface oxidation potential of human neuromelanin reveals a spherical architecture with a pheomelanin core and a eumelanin surface. Proc Natl Acad Sci U S A 103:14785–14789.

Butkovich LM, Houser MC, Chalermpalanupap T, Porter-Stransky KA, Iannitelli AF, Boles JS, Lloyd GM, Coomes AS, Eidson LN, De Sousa Rodrigues ME, Oliver DL, Kelly SD, Chang J, Bengoa-Vergniory N, Wade-Martins R, Giasson BI, Joers V, Weinshenker D, Tansey MG (2020) Transgenic Mice Expressing Human alpha-Synuclein in Noradrenergic Neurons Develop Locus Ceruleus Pathology and Nonmotor Features of Parkinson’s Disease. J Neurosci 40:7559–7576.

Carballo-Carbajal I, Laguna A, Romero-Gimenez J, Cuadros T, Bove J, Martinez-Vicente M, Parent A, Gonzalez-Sepulveda M, Penuelas N, Torra A, Rodriguez-Galvan B, Ballabio A, Hasegawa T, Bortolozzi A, Gelpi E, Vila M (2019) Brain tyrosinase overexpression implicates age-dependent neuromelanin production in Parkinson’s disease pathogenesis. Nat Commun 10:973.

Carter ME, Yizhar O, Chikahisa S, Nguyen H, Adamantidis A, Nishino S, Deisseroth K, de Lecea L (2010) Tuning arousal with optogenetic modulation of locus coeruleus neurons. Nat Neurosci 13:1526–1533.

Chalermpalanupap T, Schroeder JP, Rorabaugh JM, Liles LC, Lah JJ, Levey AI, Weinshenker D (2018) Locus Coeruleus Ablation Exacerbates Cognitive Deficits, Neuropathology, and Lethality in P301S Tau Transgenic Mice. J Neurosci 38:74–92.

Chalermpalanupap T, Kinkead B, Hu WT, Kummer MP, Hammerschmidt T, Heneka MT, Weinshenker D, Levey AI (2013) Targeting norepinephrine in mild cognitive impairment and Alzheimer’s disease. Alzheimers Res Ther 5:21.

Chalour N, Maoui A, Rat P, Massicot F, Dutot M, Faussat AM, Devevre E, Limb A, Warnet JM, Treton J, Dinet V, Mascarelli F (2018) AbetaPP-induced UPR Transcriptomic Signature of Glial Cells to Oxidative Stress as an Adaptive Mechanism to Preserve Cell Function and Survival. Curr Alzheimer Res 15:643–654.

Chen Y, Chen L, Lun ATL, Baldoni PL, Smyth GK (2025) edgeR v4: powerful differential analysis of sequencing data with expanded functionality and improved support for small counts and larger datasets. Nucleic Acids Res 53.

Clarke DJB, Marino GB, Deng EZ, Xie Z, Evangelista JE, Ma’ayan A (2024) Rummagene: massive mining of gene sets from supporting materials of biomedical research publications. Commun Biol 7:482.

Del Tredici K, Rub U, De Vos RA, Bohl JR, Braak H (2002) Where does parkinson disease pathology begin in the brain? J Neuropathol Exp Neurol 61:413–426.

Doppler CEJ, Kinnerup MB, Brune C, Farrher E, Betts M, Fedorova TD, Schaldemose JL, Knudsen K, Ismail R, Seger AD, Hansen AK, Staer K, Fink GR, Brooks DJ, Nahimi A, Borghammer P, Sommerauer M (2021a) Regional locus coeruleus degeneration is uncoupled from noradrenergic terminal loss in Parkinson’s disease. Brain 144:2732–2744.

Doppler CEJ, Kinnerup MB, Brune C, Farrher E, Betts M, Fedorova TD, Schaldemose JL, Knudsen K, Ismail R, Seger AD, Hansen AK, Staer K, Fink GR, Brooks DJ, Nahimi A, Borghammer P, Sommerauer M (2021b) Regional locus coeruleus degeneration is uncoupled from noradrenergic terminal loss in Parkinson’s disease. Brain.

Duran-Aniotz C et al. (2023) The unfolded protein response transcription factor XBP1s ameliorates Alzheimer’s disease by improving synaptic function and proteostasis. Mol Ther 31:2240–2256.

Ehrenberg AJ, Suemoto CK, Franca Resende EP, Petersen C, Leite REP, Rodriguez RD, Ferretti-Rebustini REL, You M, Oh J, Nitrini R, Pasqualucci CA, Jacob-Filho W, Kramer JH, Gatchel JR, Grinberg LT (2018) Neuropathologic Correlates of Psychiatric Symptoms in Alzheimer’s Disease. J Alzheimers Dis 66:115–126.

Ehrenberg AJ, Nguy AK, Theofilas P, Dunlop S, Suemoto CK, Di Lorenzo Alho AT, Leite RP, Diehl Rodriguez R, Mejia MB, Rub U, Farfel JM, de Lucena Ferretti-Rebustini RE, Nascimento CF, Nitrini R, Pasquallucci CA, Jacob-Filho W, Miller B, Seeley WW, Heinsen H, Grinberg LT (2017) Quantifying the accretion of hyperphosphorylated tau in the locus coeruleus and dorsal raphe nucleus: the pathological building blocks of early Alzheimer’s disease. Neuropathol Appl Neurobiol 43:393–408.

El Manaa W, Duplan E, Goiran T, Lauritzen I, Vaillant Beuchot L, Lacas-Gervais S, Morais VA, You H, Qi L, Salazar M, Ozcan U, Chami M, Checler F, Alves da Costa C (2021) Transcription- and phosphorylation-dependent control of a functional interplay between XBP1s and PINK1 governs mitophagy and potentially impacts Parkinson disease pathophysiology. Autophagy 17:4363–4385.

Falgas N, Pena-Gonzalez M, Val-Guardiola A, Perez-Millan A, Guillen N, Sarto J, Esteller D, Bosch B, Fernandez-Villullas G, Tort-Merino A, Maya G, Auge JM, Iranzo A, Balasa M, Llado A, Morales-Ruiz M, Bargallo N, Munoz-Moreno E, Grinberg LT, Sanchez-Valle R (2024) Locus coeruleus integrity and neuropsychiatric symptoms in a cohort of early- and late-onset Alzheimer’s disease. Alzheimers Dement.

Feinstein DL, Kalinin S, Braun D (2016) Causes, consequences, and cures for neuroinflammation mediated via the locus coeruleus: noradrenergic signaling system. J Neurochem 139 Suppl 2:154–178.

Francis PT, Palmer AM, Sims NR, Bowen DM, Davison AN, Esiri MM, Neary D, Snowden JS, Wilcock GK (1985) Neurochemical studies of early-onset Alzheimer’s disease. Possible influence on treatment. N Engl J Med 313:7–11.

German DC, Manaye KF, White CL, 3rd, Woodward DJ, McIntire DD, Smith WK, Kalaria RN, Mann DM (1992) Disease-specific patterns of locus coeruleus cell loss. Ann Neurol 32:667–676.

Ghosh A, Torraville SE, Mukherjee B, Walling SG, Martin GM, Harley CW, Yuan Q (2019) An experimental model of Braak’s pretangle proposal for the origin of Alzheimer’s disease: the role of locus coeruleus in early symptom development. Alzheimers Res Ther 11:59.

Gilvesy A, Husen E, Magloczky Z, Mihaly O, Hortobagyi T, Kanatani S, Heinsen H, Renier N, Hokfelt T, Mulder J, Uhlen M, Kovacs GG, Adori C (2022) Spatiotemporal characterization of cellular tau pathology in the human locus coeruleus-pericoerulear complex by three-dimensional imaging. Acta Neuropathol 144:651–676.

Giorgi FS, Biagioni F, Galgani A, Pavese N, Lazzeri G, Fornai F (2020) Locus Coeruleus Modulates Neuroinflammation in Parkinsonism and Dementia. Int J Mol Sci 21.

Hallman H, Jonsson G (1984) Monoamine neurotransmitter metabolism in microencephalic rat brain after prenatal methylazoxymethanol treatment. Brain Res Bull 13:383–389.

Han H, Cho JW, Lee S, Yun A, Kim H, Bae D, Yang S, Kim CY, Lee M, Kim E, Lee S, Kang B, Jeong D, Kim Y, Jeon HN, Jung H, Nam S, Chung M, Kim JH, Lee I (2018) TRRUST v2: an expanded reference database of human and mouse transcriptional regulatory interactions. Nucleic Acids Res 46:D380–D386.

Heneka MT, Ramanathan M, Jacobs AH, Dumitrescu-Ozimek L, Bilkei-Gorzo A, Debeir T, Sastre M, Galldiks N, Zimmer A, Hoehn M, Heiss WD, Klockgether T, Staufenbiel M (2006) Locus ceruleus degeneration promotes Alzheimer pathogenesis in amyloid precursor protein 23 transgenic mice. J Neurosci 26:1343–1354.

Hetz C, Zhang K, Kaufman RJ (2020) Mechanisms, regulation and functions of the unfolded protein response. Nat Rev Mol Cell Biol 21:421–438.

Hunsley MS, Palmiter RD (2004) Altered sleep latency and arousal regulation in mice lacking norepinephrine. Pharmacol Biochem Behav 78:765–773.

Iannitelli AF, Weinshenker D (2023) Riddles in the dark: Decoding the relationship between neuromelanin and neurodegeneration in locus coeruleus neurons. Neurosci Biobehav Rev 152:105287.

Iannitelli AF, Segal A, Pare JF, Mulvey B, Liles LC, Sloan SA, McCann KE, Dougherty JD, Smith Y, Weinshenker D (2023a) Tyrosinase-induced neuromelanin accumulation triggers rapid dysregulation and degeneration of the mouse locus coeruleus. bioRxiv.

Iannitelli AF, Kelberman MA, Lustberg DJ, Korukonda A, McCann KE, Mulvey B, Segal A, Liles LC, Sloan SA, Dougherty JD, Weinshenker D (2022) The Neurotoxin DSP-4 Dysregulates the Locus Coeruleus-Norepinephrine System and Recapitulates Molecular and Behavioral Aspects of Prodromal Neurodegenerative Disease. bioRxiv:2022.2009.2027.509797.

Iannitelli AF, Kelberman MA, Lustberg DJ, Korukonda A, McCann KE, Mulvey B, Segal A, Liles LC, Sloan SA, Dougherty JD, Weinshenker D (2023b) The Neurotoxin DSP-4 Dysregulates the Locus Coeruleus-Norepinephrine System and Recapitulates Molecular and Behavioral Aspects of Prodromal Neurodegenerative Disease. eNeuro 10.

Iversen LL, Rossor MN, Reynolds GP, Hills R, Roth M, Mountjoy CQ, Foote SL, Morrison JH, Bloom FE (1983) Loss of pigmented dopamine-beta-hydroxylase positive cells from locus coeruleus in senile dementia of Alzheimer’s type. Neurosci Lett 39:95–100.

Jacobs HIL, Becker JA, Kwong K, Engels-Dominguez N, Prokopiou PC, Papp KV, Properzi M, Hampton OL, d’Oleire Uquillas F, Sanchez JS, Rentz DM, El Fakhri G, Normandin MD, Price JC, Bennett DA, Sperling RA, Johnson KA (2021) In vivo and neuropathology data support locus coeruleus integrity as indicator of Alzheimer’s disease pathology and cognitive decline. Sci Transl Med 13:eabj2511.

Kang SS, Liu X, Ahn EH, Xiang J, Manfredsson FP, Yang X, Luo HR, Liles LC, Weinshenker D, Ye K (2020) Norepinephrine metabolite DOPEGAL activates AEP and pathological Tau aggregation in locus coeruleus. J Clin Invest 130:422–437.

Kang SS, Meng L, Zhang X, Wu Z, Mancieri A, Xie B, Liu X, Weinshenker D, Peng J, Zhang Z, Ye K (2022) Tau modification by the norepinephrine metabolite DOPEGAL stimulates its pathology and propagation. Nat Struct Mol Biol 29:292–305.

Keenan AB, Torre D, Lachmann A, Leong AK, Wojciechowicz ML, Utti V, Jagodnik KM, Kropiwnicki E, Wang Z, Ma’ayan A (2019) ChEA3: transcription factor enrichment analysis by orthogonal omics integration. Nucleic Acids Res 47:W212–W224.

Kelberman MA, Anderson CR, Chlan E, Rorabaugh JM, McCann KE, Weinshenker D (2022) Consequences of Hyperphosphorylated Tau in the Locus Coeruleus on Behavior and Cognition in a Rat Model of Alzheimer’s Disease. J Alzheimers Dis 86:1037–1059.

Kelberman MA, Rorabaugh JM, Anderson CR, Marriott A, DePuy SD, Rasmussen K, McCann KE, Weiss JM, Weinshenker D (2023) Age-dependent dysregulation of locus coeruleus firing in a transgenic rat model of Alzheimer’s disease. Neurobiol Aging 125:98–108.

Kelly L, Seifi M, Ma R, Mitchell SJ, Rudolph U, Viola KL, Klein WL, Lambert JJ, Swinny JD (2021) Identification of intraneuronal amyloid beta oligomers in locus coeruleus neurons of Alzheimer’s patients and their potential impact on inhibitory neurotransmitter receptors and neuronal excitability. Neuropathol Appl Neurobiol 47:488–505.

Korotkevich G, Sukhov V, Budin N, Shpak B, Artyomov MN, Sergushichev A (2021) Fast gene set enrichment analysis. bioRxiv.

Korukonda A et al. (2026) Pathogenic tau in the mouse locus coeruleus produces noradrenergic hyperactivity and neuropsychiatric phenotypes reminiscent of early Alzheimer’s disease. bioRxiv.

Laguna A, Penuelas N, Gonzalez-Sepulveda M, Nicolau A, Arthaud S, Guillard-Sirieix C, Lorente-Picon M, Compte J, Miquel-Rio L, Xicoy H, Liu J, Parent A, Cuadros T, Romero-Gimenez J, Pujol G, Gimenez-Llort L, Fort P, Bortolozzi A, Carballo-Carbajal I, Vila M (2024) Modelling human neuronal catecholaminergic pigmentation in rodents recapitulates age-related neurodegenerative deficits. Nat Commun 15:8819.

Li Y, Wang C, Wang J, Zhou Y, Ye F, Zhang Y, Cheng X, Huang Z, Liu K, Fei G, Zhong C, Zeng M, Jin L (2019) Mild cognitive impairment in de novo Parkinson’s disease: A neuromelanin MRI study in locus coeruleus. Mov Disord 34:884–892.

Lustberg D, Tillage RP, Bai Y, Pruitt M, Liles LC, Weinshenker D (2020) Noradrenergic circuits in the forebrain control affective responses to novelty. Psychopharmacology (Berl) 237:3337–3355.

Lustberg DJ, Liu JQ, Iannitelli AF, Vanderhoof SO, Liles LC, McCann KE, Weinshenker D (2022) Norepinephrine and dopamine contribute to distinct repetitive behaviors induced by novel odorant stress in male and female mice. Horm Behav 144:105205.

Maldonado R (1997) Participation of noradrenergic pathways in the expression of opiate withdrawal: biochemical and pharmacological evidence. Neurosci Biobehav Rev 21:91–104.

Mann DM, Yates PO (1983a) Pathological basis for neurotransmitter changes in Parkinson’s disease. Neuropathol Appl Neurobiol 9:3–19.

Mann DM, Yates PO (1983b) Possible role of neuromelanin in the pathogenesis of Parkinson’s disease. Mech Ageing Dev 21:193–203.

Mann DM, Lincoln J, Yates PO, Stamp JE, Toper S (1980) Changes in the monoamine containing neurones of the human CNS in senile dementia. Br J Psychiatry 136:533–541.

Matchett BJ, Grinberg LT, Theofilas P, Murray ME (2021) The mechanistic link between selective vulnerability of the locus coeruleus and neurodegeneration in Alzheimer’s disease. Acta Neuropathol 141:631–650.

Mathys H, Davila-Velderrain J, Peng Z, Gao F, Mohammadi S, Young JZ, Menon M, He L, Abdurrob F, Jiang X, Martorell AJ, Ransohoff RM, Hafler BP, Bennett DA, Kellis M, Tsai LH (2019) Single-cell transcriptomic analysis of Alzheimer’s disease. Nature 570:332–337.

Mathys H et al. (2023) Single-cell atlas reveals correlates of high cognitive function, dementia, and resilience to Alzheimer’s disease pathology. Cell 186:4365–4385 e4327.

Mazei-Robison MS, Nestler EJ (2012) Opiate-induced molecular and cellular plasticity of ventral tegmental area and locus coeruleus catecholamine neurons. Cold Spring Harb Perspect Med 2:a012070.

McCall A, Forouhandehpour R, Celebi S, Richard-Malenfant C, Hamati R, Guimond S, Tuominen L, Weinshenker D, Jaworska N, McQuaid RJ, Shlik J, Robillard R, Kaminsky Z, Cassidy CM (2024) Evidence for Locus Coeruleus-Norepinephrine System Abnormality in Military Posttraumatic Stress Disorder Revealed by Neuromelanin-Sensitive Magnetic Resonance Imaging. Biol Psychiatry 96:268–277.

Mercan D, Heneka MT (2022) The Contribution of the Locus Coeruleus-Noradrenaline System Degeneration during the Progression of Alzheimer’s Disease. Biology (Basel) 11.

Meyer C et al. (2025) Early Locus Coeruleus noradrenergic axon loss drives olfactory dysfunction in Alzheimer’s disease. Nat Commun 16:7338.

Mulvey B, Frye HE, Lintz T, Fass S, Tycksen E, Nelson EC, Moron JA, Dougherty JD (2023) Cnih3 Deletion Dysregulates Dorsal Hippocampal Transcription across the Estrous Cycle. eNeuro 10.

Mulvey B, Bhatti DL, Gyawali S, Lake AM, Kriaucionis S, Ford CP, Bruchas MR, Heintz N, Dougherty JD (2018) Molecular and Functional Sex Differences of Noradrenergic Neurons in the Mouse Locus Coeruleus. Cell Rep 23:2225–2235.

Murchison CF, Zhang XY, Zhang WP, Ouyang M, Lee A, Thomas SA (2004) A distinct role for norepinephrine in memory retrieval. Cell 117:131–143.

Naegeli C, Zeffiro T, Piccirelli M, Jaillard A, Weilenmann A, Hassanpour K, Schick M, Rufer M, Orr SP, Mueller-Pfeiffer C (2018) Locus Coeruleus Activity Mediates Hyperresponsiveness in Posttraumatic Stress Disorder. Biol Psychiatry 83:254–262.

Pletnikova O, Kageyama Y, Rudow G, LaClair KD, Albert M, Crain BJ, Tian J, Fowler D, Troncoso JC (2018) The spectrum of preclinical Alzheimer’s disease pathology and its modulation by ApoE genotype. Neurobiol Aging 71:72–80.

Poe GR, Foote S, Eschenko O, Johansen JP, Bouret S, Aston-Jones G, Harley CW, Manahan-Vaughan D, Weinshenker D, Valentino R, Berridge C, Chandler DJ, Waterhouse B, Sara SJ (2020) Locus coeruleus: a new look at the blue spot. Nat Rev Neurosci 21:644–659.

Porter-Stransky KA, Centanni SW, Karne SL, Odil LM, Fekir S, Wong JC, Jerome C, Mitchell HA, Escayg A, Pedersen NP, Winder DG, Mitrano DA, Weinshenker D (2019) Noradrenergic Transmission at Alpha1-Adrenergic Receptors in the Ventral Periaqueductal Gray Modulates Arousal. Biol Psychiatry 85:237–247.

Prediger RD, Matheus FC, Schwarzbold ML, Lima MM, Vital MA (2012) Anxiety in Parkinson’s disease: a critical review of experimental and clinical studies. Neuropharmacology 62:115–124.

Prokopiou PC, Engels-Dominguez N, Papp KV, Scott MR, Schultz AP, Schneider C, Farrell ME, Buckley RF, Quiroz YT, El Fakhri G, Rentz DM, Sperling RA, Johnson KA, Jacobs HIL (2022) Lower novelty-related locus coeruleus function is associated with Abeta-related cognitive decline in clinically healthy individuals. Nat Commun 13:1571.

Ranjbar-Slamloo Y, Fazlali Z (2019) Dopamine and Noradrenaline in the Brain; Overlapping or Dissociate Functions? Front Mol Neurosci 12:334.

Raskind MA, Peskind ER, Holmes C, Goldstein DS (1999) Patterns of cerebrospinal fluid catechols support increased central noradrenergic responsiveness in aging and Alzheimer’s disease. Biol Psychiatry 46:756–765.

Remy P, Doder M, Lees A, Turjanski N, Brooks D (2005) Depression in Parkinson’s disease: loss of dopamine and noradrenaline innervation in the limbic system. Brain 128:1314–1322.

Rorabaugh JM, Chalermpalanupap T, Botz-Zapp CA, Fu VM, Lembeck NA, Cohen RM, Weinshenker D (2017) Chemogenetic locus coeruleus activation restores reversal learning in a rat model of Alzheimer’s disease. Brain 140:3023–3038.

Smyth GK, Michaud J, Scott HS (2005) Use of within-array replicate spots for assessing differential expression in microarray experiments. Bioinformatics 21:2067–2075.

Sommerauer M, Fedorova TD, Hansen AK, Knudsen K, Otto M, Jeppesen J, Frederiksen Y, Blicher JU, Geday J, Nahimi A, Damholdt MF, Brooks DJ, Borghammer P (2018) Evaluation of the noradrenergic system in Parkinson’s disease: an 11C-MeNER PET and neuromelanin MRI study. Brain 141:496–504.

Subramanian A, Tamayo P, Mootha VK, Mukherjee S, Ebert BL, Gillette MA, Paulovich A, Pomeroy SL, Golub TR, Lander ES, Mesirov JP (2005) Gene set enrichment analysis: a knowledge-based approach for interpreting genome-wide expression profiles. Proc Natl Acad Sci U S A 102:15545–15550.

Sulzer D, Mosharov E, Talloczy Z, Zucca FA, Simon JD, Zecca L (2008) Neuronal pigmented autophagic vacuoles: lipofuscin, neuromelanin, and ceroid as macroautophagic responses during aging and disease. J Neurochem 106:24–36.

Sulzer D, Bogulavsky J, Larsen KE, Behr G, Karatekin E, Kleinman MH, Turro N, Krantz D, Edwards RH, Greene LA, Zecca L (2000) Neuromelanin biosynthesis is driven by excess cytosolic catecholamines not accumulated by synaptic vesicles. Proc Natl Acad Sci U S A 97:11869–11874.

Tansey MG, Wallings RL, Houser MC, Herrick MK, Keating CE, Joers V (2022) Inflammation and immune dysfunction in Parkinson disease. Nat Rev Immunol 22:657–673.

Theofilas P, Ehrenberg AJ, Dunlop S, Di Lorenzo Alho AT, Nguy A, Leite REP, Rodriguez RD, Mejia MB, Suemoto CK, Ferretti-Rebustini REL, Polichiso L, Nascimento CF, Seeley WW, Nitrini R, Pasqualucci CA, Jacob Filho W, Rueb U, Neuhaus J, Heinsen H, Grinberg LT (2017) Locus coeruleus volume and cell population changes during Alzheimer’s disease progression: A stereological study in human postmortem brains with potential implication for early-stage biomarker discovery. Alzheimers Dement 13:236–246.

Tillage RP, Wilson GE, Liles LC, Holmes PV, Weinshenker D (2020a) Chronic Environmental or Genetic Elevation of Galanin in Noradrenergic Neurons Confers Stress Resilience in Mice. J Neurosci 40:7464–7474.

Tillage RP, Sciolino NR, Plummer NW, Lustberg D, Liles LC, Hsiang M, Powell JM, Smith KG, Jensen P, Weinshenker D (2020b) Elimination of galanin synthesis in noradrenergic neurons reduces galanin in select brain areas and promotes active coping behaviors. Brain Struct Funct 225:785–803.

Vazey EM, Aston-Jones G (2012) The emerging role of norepinephrine in cognitive dysfunctions of Parkinson’s disease. Front Behav Neurosci 6:48.

Wang ZM, Grinevich V, Meeker WR, Zhang J, Messi ML, Budygin E, Delbono O (2025) Early signs of neuron autonomous and non-autonomous hyperexcitability in locus coeruleus noradrenergic neurons of a mouse model of tauopathy and Alzheimer’s disease. Acta Physiol (Oxf) 241:e70022.

Weinshenker D (2018) Long Road to Ruin: Noradrenergic Dysfunction in Neurodegenerative Disease. Trends Neurosci 41:211–223.

Williams JT, North RA, Shefner SA, Nishi S, Egan TM (1984) Membrane properties of rat locus coeruleus neurones. Neuroscience 13:137–156.

Xie Z, Bailey A, Kuleshov MV, Clarke DJB, Evangelista JE, Jenkins SL, Lachmann A, Wojciechowicz ML, Kropiwnicki E, Jagodnik KM, Jeon M, Ma’ayan A (2021) Gene Set Knowledge Discovery with Enrichr. Curr Protoc 1:e90.

Ye R, O’Callaghan C, Rua C, Hezemans FH, Holland N, Malpetti M, Jones PS, Barker RA, Williams-Gray CH, Robbins TW, Passamonti L, Rowe J (2022) Locus Coeruleus Integrity from 7 T MRI Relates to Apathy and Cognition in Parkinsonian Disorders. Mov Disord 37:1663–1672.

Zarow C, Lyness SA, Mortimer JA, Chui HC (2003) Neuronal loss is greater in the locus coeruleus than nucleus basalis and substantia nigra in Alzheimer and Parkinson diseases. Arch Neurol 60:337–341.

Zecca L, Zucca FA, Wilms H, Sulzer D (2003) Neuromelanin of the substantia nigra: a neuronal black hole with protective and toxic characteristics. Trends Neurosci 26:578–580.

Zecca L, Wilms H, Geick S, Claasen JH, Brandenburg LO, Holzknecht C, Panizza ML, Zucca FA, Deuschl G, Sievers J, Lucius R (2008a) Human neuromelanin induces neuroinflammation and neurodegeneration in the rat substantia nigra: implications for Parkinson’s disease. Acta Neuropathol 116:47–55.

Zecca L, Stroppolo A, Gatti A, Tampellini D, Toscani M, Gallorini M, Giaveri G, Arosio P, Santambrogio P, Fariello RG, Karatekin E, Kleinman MH, Turro N, Hornykiewicz O, Zucca FA (2004) The role of iron and copper molecules in the neuronal vulnerability of locus coeruleus and substantia nigra during aging. Proc Natl Acad Sci U S A 101:9843–9848.

Zecca L, Bellei C, Costi P, Albertini A, Monzani E, Casella L, Gallorini M, Bergamaschi L, Moscatelli A, Turro NJ, Eisner M, Crippa PR, Ito S, Wakamatsu K, Bush WD, Ward WC, Simon JD, Zucca FA (2008b) New melanic pigments in the human brain that accumulate in aging and block environmental toxic metals. Proc Natl Acad Sci U S A 105:17567–17572.

Zhang W, Phillips K, Wielgus AR, Liu J, Albertini A, Zucca FA, Faust R, Qian SY, Miller DS, Chignell CF, Wilson B, Jackson-Lewis V, Przedborski S, Joset D, Loike J, Hong JS, Sulzer D, Zecca L (2011) Neuromelanin activates microglia and induces degeneration of dopaminergic neurons: implications for progression of Parkinson’s disease. Neurotox Res 19:63–72.

Zweig RM, Cardillo JE, Cohen M, Giere S, Hedreen JC (1993) The locus ceruleus and dementia in Parkinson’s disease. Neurology 43:986–991.

