## Supplemental Table 1 for "Tyrosinase-induced neuromelanin accumulation triggers rapid dysregulation and degeneration of the mouse locus coeruleus"

**Table 1. IHC antibodies**

| **Antibodies** | **Host** | **Manufacturer** | **Catalog #** | **Dilution** |
| --- | --- | --- | --- | --- |
| Tyrosine hydroxylase | Chicken | Abcam | ab76442 | 1:1000 |
| Norepinephrine transporter | Mouse | Mab Technologies | NET05-2 | 1:1000 |
| Human tyrosinase | Mouse | Thermo Fisher Scientific | MS-800-B1 | 1:500 |
| GFAP | Rabbit | Abcam | AB7260 | 1:1000 |
| GFAP | Guinea Pig | Synaptic Systems | 173 004 | 1:1000 |
| Iba1 | Rabbit | FujiFilm | 019-19741 | 1:1000 |
| NeuroTrace 435/455 Blue Fluorescent Nissl Stain | NA | Thermo Fisher Scientific | N21479 | 1:500 |
| NeuroTrace 640/660 Far Red Fluorescent Nissl Stain | N/A | Thermo Fisher Scientific | N21483 | 1:250 |
| Alexa Fluor 488 Anti-Chicken | Goat | Abcam | ab150169 | 1:500 |
| Alexa Fluor 488 Anti-Rabbit | Goat | Thermo Fisher Scientific | A-11008 | 1:500 |
| Alexa Fluor 488 Anti-Guinea Pig | Goat | Thermo Fisher Scientific | A-11073 | 1:500 |
| Alexa Fluor 568 Anti-Rabbit | Goat | Thermo Fisher Scientific | A-11011 | 1:500 |
| Alexa Fluor 568 Anti-Chicken | Goat | Thermo Fisher Scientific | A-11041 | 1:500 |
| Alexa Fluor 568 Anti-Rabbit | Goat | Thermo Fisher Scientific | A-11011 | 1:500 |
