## Supplemental Table 2 for "Tyrosinase-induced neuromelanin accumulation triggers rapid dysregulation and degeneration of the mouse locus coeruleus"

**Supplemental Table 2. Monoamine levels measured by HPLC at 1 week.** Data shown as mean ± SEM. N=5-6 per group. *****p<0.05, **p<0/01.

|  | **Pons** | | **PFC** | | **Hippocampus** | |
| --- | --- | --- | --- | --- | --- | --- |
|  | **EYFP** | **hTyr** | **EYFP** | **hTyr** | **EYFP** | **hTyr** |
| **DA** | 11.37 ± 0.69 | 10.39 ± 0.91 | 20.14 ± 8.32 | 4.54 ± 0.38 | 3.53 ± 1.29 | 2.33 ± 0.28 |
| **DOPAC** | 6.98 ± 0.49 | 7.53 ± 0.72 | 3.08 ± 0.61 | 2.40 ± 0.20 | 2.52 ± 0.26 | 2.47 ± 0.45 |
| **DOPAC:DA** | 0.62 ± 0.03 | 0.72 ± 0.02** | 0.30 ± 0.08 | 0.53 ± 0.02* | 1.04 ± 0.20 | 1.11 ± 0.19 |
| **5-HT** | 170.9 ± 3.38 | 162.5 ± 4.33 | 40.66 ± 3.34 | 45.30 ± 2.94 | 69.17 ± 3.43 | 63.04 ± 3.73 |
| **5-HIAA** | 58.31 ± 3.09 | 61.95 ± 4.11 | 7.72 ± 0.22 | 8.59 ± 0.45 | 7.72 ± 0.22 | 8.59 ± 0.45 |
| **5-HIAA:5-HT** | 0.34 ± 0.02 | 0.38 ± 0.20 | 0.20 ± 0.02 | 0.19 ± 0.01 | 0.36 ± 0.03 | 0.38 ± 0.01 |
